# Sequence determinants of efficient exon 44 skipping in Duchenne muscular dystrophy define design principles for steric-blocking antisense oligonucleotides

**DOI:** 10.64898/2026.06.29.735365

**Authors:** Enyang Han, Katherine Webster, Tiberiu Loredan Stan, Christa Tanganyika-de Winter, Elizabeth van der Pijl, Jorden Tahquechi, Brian Heglar, Chad Koehler, Irinna Papangeli, Donnie Mackenzie, Brett E. Crawford, Annemieke Aartsma-Rus, Tom A. Hartl

**Affiliations:** BioMarin Pharmaceutical Inc., Novato, CA, USA; Department of Human Genetics, Leiden University Medical Center (LUMC), Leiden, The Netherlands

**Keywords:** splice-switching oligonucleotide, steric-blocking oligonucleotide, gymnotic delivery, exonic splicing enhancer, humanized mouse model, Becker muscular dystrophy, nucleofection

## Abstract

Duchenne muscular dystrophy (DMD) is caused by mutations in the *DMD* gene that disrupt the reading frame and abolish expression of functional dystrophin protein. Antisense oligonucleotides (ASO) can restore production of partially functional dystrophins by inducing exon skipping to restore the reading frame of dystrophin transcripts. While exon skipping is an FDA approved therapeutic strategy, there are currently no approved therapies for patients amenable to exon 44 skipping (8% of DMD patients). Here, we carried out a discovery campaign to identify phosphorothioate (PS) ASOs that efficiently induce exon 44 skipping and to define key sequence and chemistry features associated with activity. A tiling and micro-tiling approach with 18mer fully PS and 2’-O-methoxyethyl (2’MOE) modified ASOs in patient-derived myotubes identified five exonic target regions that promote skipping. ASO activity was strongly correlated across skeletal muscle and iPSC-derived cardiomyocytes, indicating similar exon 44 splicing regulation across cell types. Optimization studies showed that for 2’MOE PS ASOs, 16–20mers were generally most active, while longer ASOs often had reduced activity due in part to impaired productive uptake into cells. Swapping out 2’MOE modifications at both terminal positions for locked nucleic acids (LNAs) rarely improved activity and could also reduce it. Finally, promising candidates were tested in a humanized mouse model with an exon 44 skippable deletion, where one ASO outperformed others, inducing dose-dependent exon 44 skipping and dystrophin restoration in mouse models. These findings define practical design rules for exon 44-targeted ASOs and provide a foundation for therapeutic development.

## Introduction

Duchenne muscular dystrophy (DMD) is a severe, X-linked disorder caused by mutations in the *DMD* gene that abolish expression of functional dystrophin, a protein essential for maintaining sarcolemmal stability during muscle contraction. Loss of dystrophin leads to progressive muscle fiber degeneration, chronic inflammation, and replacement of muscle with fibrotic and adipose tissue, ultimately affecting skeletal, cardiac, and respiratory muscle function [1]. Clinically, DMD presents in early childhood with delayed motor milestones, progressive proximal muscle weakness, and loss of ambulation, followed by cardiomyopathy and respiratory insufficiency [1]. A key determinant of disease severity is the “reading frame rule,” whereby out-of-frame mutations result in absence of functional dystrophin and a DMD phenotype, while in-frame mutations permit production of a truncated but partially functional protein and lead to the milder Becker muscular dystrophy (BMD) phenotype [2–4]. This genotype–phenotype relationship provides the mechanistic foundation for therapeutic strategies aimed at restoring the translational reading frame.

Antisense oligonucleotide (ASO)-mediated exon skipping is a mutation-specific therapeutic strategy that exploits the reading frame rule by redirecting pre-mRNA splicing to restore the open reading frame of *DMD* transcripts. Targeted skipping of specific exons can reframe transcripts and enable production of a truncated but functional dystrophin protein analogous to that observed in BMD patients [5, 6].

Currently, the FDA has approved 4 ASOs targeting 3 dystrophin exons via the accelerated approval pathway: eteplirsen (exon 51 skipping), golodirsen and viltolarsen (exon 53 skipping), and casimersen (exon 45 skipping). Approval was based on the ASO’s ability to restore low levels of dystrophin expression in muscle [7–10]. These therapies are applicable to defined subsets of patients, with approximately ~13% of DMD patients amenable to exon 51 skipping, ~10% to exon 53 skipping, and ~8% to exon 45 skipping. As 2% of patients carry a deletion of exon 52, which can be corrected either by exon 51 or exon 53 skipping, collectively the approved ASOs cover roughly 25–30% of the DMD population [11, 12]. While these approvals validate exon skipping as a therapeutic approach, the levels of dystrophin restoration achieved remain modest and clinical benefit is still being clarified, motivating continued innovation in ASO chemistry, delivery, and target selection. Current approaches in clinical and preclinical development include next-generation chemistries with improved potency and tissue delivery, peptide-conjugated PMOs, and antibody-ASO-conjugate strategies designed to expand applicability and enhance functional outcomes [13]. Together, these efforts aim to overcome the limitations of first-generation exon-skipping therapies and improve therapeutic efficacy across a broader range of DMD mutations.

While multiple exon 44 skipping approaches are in clinical development, including programs from NS Pharma, Avidity Biosciences, and Entrada Therapeutics, there are no approved antisense oligonucleotide (ASO) therapies for this patient population, which makes up 8% of DMD [11, 13–15]. Here, we carried out a discovery campaign to identify regions within DMD exon 44 that drive efficient exon skipping when targeted by phosphorothioate (PS) ASOs with full 2’MOE modifications in muscle and cardiac cells. Remarkably, ASO potency was highly concordant between muscle and cardiac cell types, suggesting shared regulatory mechanisms governing exon 44 splicing. We found that optimal ASO lengths ranged between 16 and 20 nts, consistent with what has been observed by others [16]. Shorter ASOs likely suffer from reduced binding affinity, whereas longer ASOs may exhibit diminished cellular uptake. Interestingly, incorporation of terminal locked nucleic acids (LNAs) to enhance binding affinity rarely improved activity and could also decrease exon skipping activity relative to non-LNA ASOs, consistent with prior reports that excessive affinity or altered protein interactions can negatively impact ASO efficacy [17, 18]. These findings provide a framework for exon 44-targeted therapeutic development and may be broadly applicable to other steric-blocking ASO programs.

## Methods

### ASOs

ASO sequences are given in Table S1. ASOs were ordered with 2’-O-methoxyethyl (MOE) RNA and/or locked nucleic acid (LNA) nucleotides (nts) and a full length phosphorothioate (PS) backbone. ASOs were produced by IDT and diluted to 500 µM in 1X IDTE buffer (10 mM Tris, 0.1 mM EDTA, pH 8) and stored at −20 degrees C until further use.

### Gymnotic cellular assays

DMD patient-derived myoblasts carrying either an exon 42–43 deletion (7796) or an exon 52 deletion (KM571) were obtained from the Institut de Myologie and had been previously immortalized using hTERT and CDK4 [19]. Myoblasts were maintained in Skeletal Muscle Cell Growth Medium (PromoCell, Cat. No. C-23060) supplemented according to the manufacturer’s instructions, including 12.5% heat-inactivated fetal bovine serum (Invitrogen, Cat. No. 10270-106) and 100 U/mL penicillin-streptomycin (Invitrogen, cat#: 15140-122). Cells were plated at 25,000 cells per well in collagen I-coated 96-well plates (Corning Cat. No. 354649).

After 48 h, differentiation was initiated by replacing growth medium with differentiation medium consisting of low-glucose DMEM containing pyruvate and lacking glutamine and phenol red (Thermo Fisher Scientific, Cat. No. 11054-020), supplemented with 2% heat-inactivated fetal bovine serum (Invitrogen, Cat. No. 10270-106), 1% glucose (45% w/v; Sigma-Aldrich, Cat. No. G8769), 2% GlutaMAX Supplement (Invitrogen, Cat. No. 35050-038), and 1% penicillin-streptomycin (Invitrogen, Cat. No. 15140-122). Cells were differentiated for 7 days to generate myotubes and then they were treated with ASOs at 1 μM or 4 μM by gymnotic uptake for 72 h. Total RNA was isolated using the Cells-to-CT kit (Thermo Fisher Scientific; 55 μL final lysis volume), and 10 μL of lysate was reverse transcribed according to the manufacturer’s instructions. Exon 44 skipping was quantified by TaqMan qPCR using 4 μL of cDNA per reaction. Skipped transcripts were detected using the following primers in exons 41 and 45 and a probe spanning the Exon 41/45 junction. Expression was normalized to UBC using a commercial TaqMan assay (IDT; Hs.PT.39a.22214853), and relative exon skipping was calculated using the ΔΔCt method. To help control for plate-to-plate variability in this primary screen, each plate carried a negative control (vehicle) and a positive control ASO (DMD44 (27, 46)).

### Nucleofection cellular assays

On day 1, 100,000 myoblasts per well were nucleofected (Lonza, setting DS-150, SF solution) in the presence of 1.4 μM of the individual oligonucleotides, diluted 10-fold with Skeletal Muscle growth medium (PromoCell (cat#: C-23060), final volume 12.5% HI-FBS and 100 units/mL Pen/Strep), and then transferred to a 96-well tissue culture plate. On day 4, the cells were harvested using TaqMan Gene Expression Cells-to-CT (Thermo Fisher Scientific, 55 microliter final lysis volume), 10 μL of RNA was reverse transcribed, and then 4 μL of the resulting cDNA were evaluated by qPCR.

### *In vivo* studies in hDMD mice

Initial in vivo studies were carried out with hDMD mice in a wildtype mouse dystrophin background as a model system to assess exon 44 skipping [20]. *In vivo* studies were performed in accordance with proper IACUC standards taken from The Guide and underwent IACUC peer review. Five to eight mice aged between 8-10 weeks were administered ASO at 1.71 μmol/kg or 3.42 μmol/kg intravenously via lateral tail vein injection once weekly for 4 consecutive weeks. ASOs were obtained from AxoLabs. Mice were sacrificed one week after the 4th dose and tissues were and flash frozen. RNA was extracted from frozen samples by tissue pulverization using a Covaris Cryoprep and further homogenized in Zymo RNA/DNA Shield buffer using an Omni Bead Ruptor. The RT-PCR reactions used 150-200 ng of RNA to generate cDNA with Cells-to-CT Bulk RT Reagents kit (ThermoFisher) which then was analyzed by ddPCR for exon 44 skipping with primer sets that enable copy number quantification for the non-skipped exon junction, the skipped exon 43/45 junction, and mouse UBC as a housekeeping gene.

### *In vivo* studies in DMDdel45/*mdx* mice

A hDMDdel45/*mdx* mouse was implemented at Leiden University Medical Center (LUMC) to evaluate exon 44 skipping. The hDMDdel45/*mdx* mouse model enables preclinical testing of antisense oligonucleotides (AONs) designed to induce skipping of human DMD exon 44. It carries mutated versions of both the murine and human DMD genes, so it does not produce either mouse or human dystrophin. This makes it suitable for pharmacokinetic and pharmacodynamic studies of AONs targeting human DMD exons within a dystrophic context [21]. As a result, findings can be translated more directly to clinical settings, minimizing the need to extrapolate from mouse-specific to human-specific AONs. The hDMDdel45/*mdx* mice were bred at the LUMC animal facility. They were housed in individually ventilated cages maintained at 20.5 °C under a 12-hour light/dark cycle, with free access to water and standard RM3 chow throughout the study. All procedures were approved by the LUMC Animal Welfare Body (ethical permit PE.17306.02.005).

To test the efficiency of the AONs to target muscle cells and induce exon skipping, the compounds were intravenously injected, weekly, in blinded, randomized cohorts of 5 male mice per treatment group, as follows:

1. hDMDdel45/*mdx* + saline (n=5)
2. hDMDdel45/*mdx* + Wdolg2053 (25.01 mg/kg) (n=5)
3. hDMDdel45/*mdx* + Wdolg2053 (125.06 mg/kg) (n=5)
4. hDMDdel45/*mdx* + Wdolg503 (22.45 mg/kg) (n=5)
5. hDMDdel45/*mdx* + Wdolg503 (4.49 mg/kg) (n=5)

Mice were 5 weeks old at the time of first injection. Animals were scheduled to receive a weekly intravenous injection, for four consecutive weeks of the above-mentioned compounds with the maximal injection volume of 150 μL single bolus and approximately 30 sec administration time /compound by an experienced technician.

One control group of hDMDdel45/*mdx* was also included, receiving weekly intravenous injections of 150 µL saline. No dystrophin restoration was expected for the hDMDdel45/*mdx* saline-injected mice. Blinding was implemented to minimize bias during experimental procedures and data collection. Specifically, the experimenters responsible for administering the injections were blinded to the identity of the compounds; each substance was labeled generically to prevent any knowledge of its contents. This ensured that treatment administration was conducted without preconceived expectations or influence. Bodyweight was recorded once a week for the duration of the experiment. A control group consisting of one hDMD/*mdx* mouse, which express human dystrophin and are therefore non-dystrophic, was included as a control group to provide control tissue.

One week after last injection animals were scheduled for sacrificing and tissue collection. In brief, animals were sedated with isoflurane (3-4% induction, ˃4% maintenance) and 5 minutes prior opening the thorax, animals were administrated few drops of subcutaneous lidocaine. Blood was collected by cardiac puncture. Serum and residuals were stored in −80 freezer.

After sacrifice, gastrocnemius, quadriceps, diaphragm, heart, liver, kidney, spleen, were isolated for further analysis.

### RT-PCR analysis for exon skip assessment

Frozen muscle tissues were homogenized in zirconium bead pre-filled tubes (1.4 mm Zirconium Beads Pre-Filled TubePFAW 1400-100-1OPS Diagnostics) containing Trisure. Homogenization was carried out using a MagNA Lyser (Roche Diagnostics). Total RNA was then extracted according to the TriSure protocol. RNA concentration was determined using a NanoDrop spectrophotometer (Thermo Fisher Scientific, Waltham, MA), and samples were stored at −80 °C.

Total RNA (1000 ng) was mixed with random hexamers (40 ng/µL) and dNTPs (10 mM). Reverse transcription was performed using 5× reaction buffer, rRNasin (40 U/µL), and Bioscript Reverse Transcriptase (200 U/µL) at 25 °C for 10 min, 42 °C for 1 h, and 70 °C for 10 min. cDNA was stored at −20 °C or used directly.

A PCR was performed in 50 µL, containing 1.5 µL cDNA, 10× SuperTaq buffer, dNTPs (10 mM), primers h43f2 and h47r2 (10 pmol/µL), and Taq DNA polymerase (5 U/µL). Cycling conditions were 94 °C for 5 min; 36 cycles of 94 °C (30 s), 60 °C (30 s), 72 °C (30 s); final extension at 72 °C for 7 min. PCR products were separated on 1.5% agarose gels and quantified by Lab on a chip.

### Dystrophin quantification

Dystrophin levels were quantified using a capillary-based Western immunoassay as previously described and validated [22, 23].

In brief, to generate protein lysates frozen muscle tissues were homogenized in zirconium bead pre-filled tubes (1.4 mm Zirconium Beads Pre-Filled TubePFAW 1400-100-1OPS Diagnostics) containing protein isolation buffer (25% SDS, 1.25 M Tris-HCl) using the MagNA Lyser (Roche). After repeated homogenization and centrifugation steps, supernatants were transferred to clean 1.5 mL tubes and heated at 95 °C for 10 min. Protein concentration was determined using a Bicinchoninic Acid (BCA) assay (Pierce™ BCA Protein Assay Kit - 1L23225, Thermo Fisher Scientific), and aliquots were stored at −80 °C. Lysates were diluted 1:10 by mixing 4 µL of protein lysate with 36 µL of protein isolation buffer, and 10 µL of this dilution was used for analysis.

Alternatively, analysis was performed on the Jess System (Bio-Techne) according to the manufacturer’s instructions. A 66–440 kDa separation module (Bio-Techne) was used in combination with the anti-rabbit Detection Module (Bio-Techne).

Dystrophin was detected using a 1:100 dilution of rabbit monoclonal antibody (ab154168, Abcam). To control for loading variability, vinculin was measured using a 1:100 dilution of anti-vinculin rabbit monoclonal antibody (42H89L44, Cell Signaling). Detection was performed with an anti-rabbit secondary antibody (Bio-Techne). Tissue samples were diluted in sample buffer (100× diluted “10× Sample Buffer 2” from the separation module) to a concentration of 1 mg/mL (2.5 µL loaded per sample), mixed with fluorescent master mix, and heated at 95 °C for 5 minutes. Samples, blocking reagent (antibody diluent), primary antibody, HRP-conjugated secondary antibody, and chemiluminescent substrate were loaded into the separation module, and runs were performed using default instrument settings.

For quantification, relative dystrophin levels were determined using a four-point calibration curve generated from untreated hDMD/*mdx* and hDMDdel45/*mdx* muscle samples. The average signals from hDMD/*mdx* mouse and hDMDdel45/*mdx*-saline controls (representing 100% and 0% dystrophin, respectively) were used to establish a trendline. Dystrophin levels were normalized to vinculin by dividing the dystrophin signal by the vinculin signal, and results were expressed as a percentage of hDMD/*mdx* mouse levels (% hDMD/*mdx* mouse) based on the calibration curve.

## Results

### Five regions on DMD Exon 44 can be targeted by ASOs to induce efficient exon skipping

To identify regions that lead to exon 44 skipping when bound by an ASO, a gymnotic cellular assay in DMD patient-derived myotubes carrying a deletion of exon 42-43 was established. It was reasoned that this approach had a high probability of identifying ASOs which could translate to *in vivo* efficacy due to it physiological and genetic relevance. When exon 44 is skipped in this genetic background the reading frame of the mature transcript is restored from out-of-frame to in-frame.

Forty-seven ASOs of 18 nts in length were designed to walk across the exon 44 region in 5 nt steps starting 50 nts upstream of the exon 44 3’ splice acceptor in intron 43 and ending 49 nts downstream of the Exon 44 5’ splice donor in intron 44. The ASOs had full PS backbones and each 2’ position of the ribose carried a MOE modification. This is the same design as the ASO Nusinersen which is FDA approved for Spinal Muscular Atrophy [24].

Thirty-nine of the 47 ASOs (83%) induced exon 44 skipping above background levels observed for the vehicle treated cells. This is consistent with prior observations that most ASOs targeting exon 44 will induce some level of skipping [25]. ASOs binding intronic regions were less likely to induce skipping compared to those targeting the exon as was also reported before [26]. There was one exception, DMD44 (−21, −4), which induced skipping at levels of about 50% of the positive control. However, because patient deletion mutations are rarely mapped at nt resolution beyond the exons deleted, this intronic ASO-binding region was not pursued due to the possibility that it may be absent in some exon 44 skip amenable patients. Five regions were pursued further as they passed our criteria as being exonic and the ASOs binding them were at least 50% the activity of the positive control ASO. Region #1 spans positions −6 to 17 (23 nts), region #2 spans positions 10-47 (38 nts), region #3 spans positions 50-82 (33 nts), region #4 spans positions 75-112 (38 nts), and region #5 spans 105-127 (23 nts).

### Further finetuning to optimize ASO target regions

To identify optimal 18mer ASOs within each region, additional sequences were designed to expand coverage and fill gaps between those tested in Fig. 1. All 18mers targeting broadened regions 1 (−10 to 21), 2 (6 to 51), 3 (46 to 86), 4 (75 to 116), and 5 (101 to 131) were evaluated in a gymnotic assay in Del42,43 myotubes. In region #1, the newly tested DMD44 (−4, 14) was about 2 times better than primary screen hits DMD44 (−6, 12) and DMD44 (−1, 17) (Fig. 2A). Microtiling across region 2 revealed two peaks of activity that were not apparent from the 5 bp walk in Fig. 1. Five better ASOs were uncovered from the single nt walk compared to the hits from the primary screen (Fig. 2B). No newly tested ASOs in regions 3 and 5 were better than those found in the primary screen (Figs. 2C, 2E). While no clearly better ASOs were found from microtiling region #4, several others equipotent to the primary hits were discovered. Overall targeting regions #4 and #2 appeared to be the best way to induce exon 44 skipping followed by region #1, then targeting regions #3 and #5 induced the least skipping. The top ASO in region #1 is DMD44 (−4, 14). Region #2’s top ASOs are DMD44 (13, 30) in the first peak and DMD (29, 46) in the second peak. Given that the top ASOs for region #2’s two peaks overlap by only 0 to 2 nts, and the ASOs between them have less activity, it is likely that the two peaks represent different splice enhancer motifs. Region #4’s top ranked ASO is DMD44 (97, 114).

**Figure 1.**
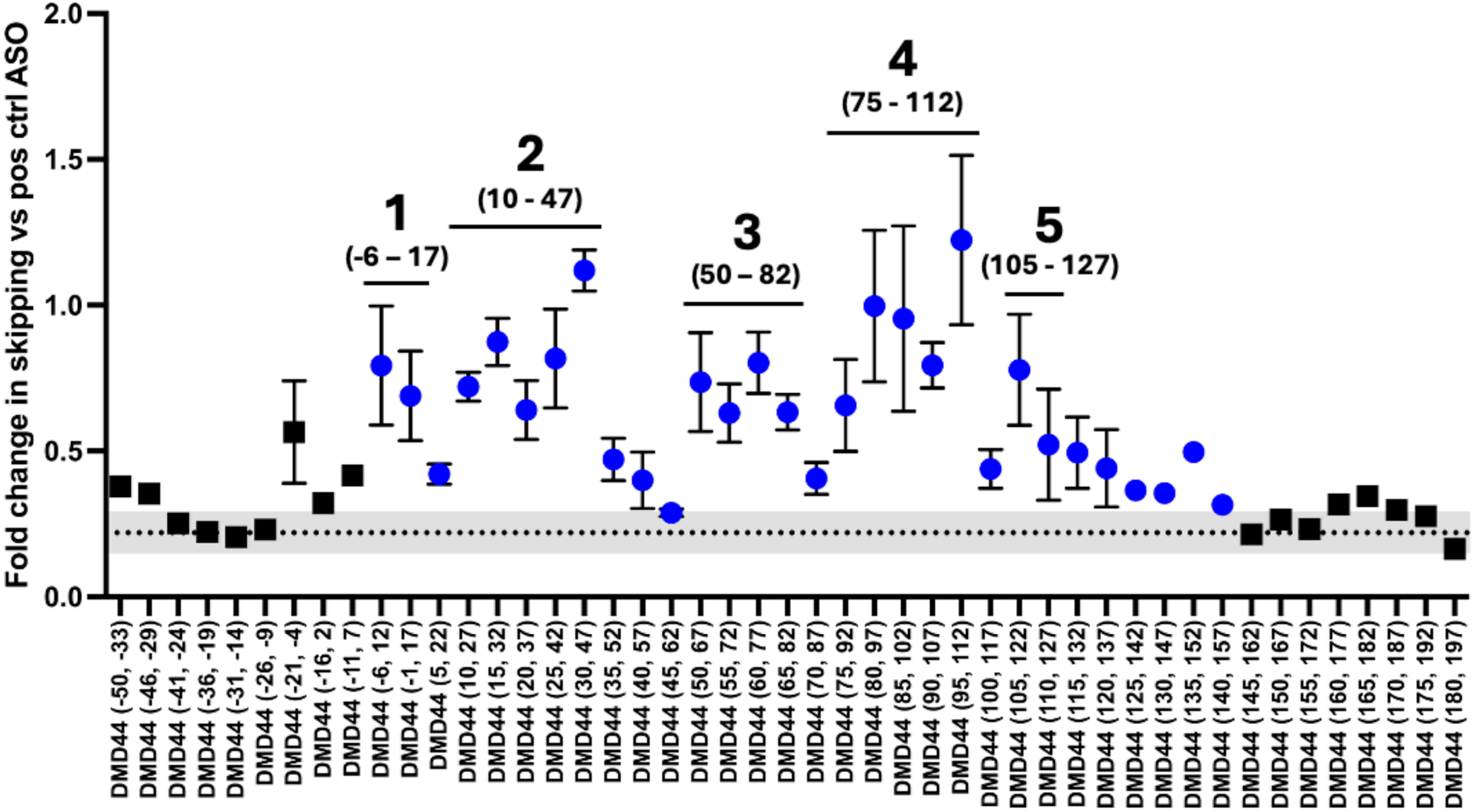
Primary ASO screen identified 5 regions on DMD Exon 44 that lead to efficient exon skipping when targeted by ASOs. Forty-seven ASOs were designed to walk in 5 nt steps across the exon 44 region starting 50 nts in intron 43 and ending 49 nts into intron 45. Data on the Y axis are expressed as fold change relative to a positive control ASO (DMD44 (27, 46)). ASOs bound to ≥50% intronic sequence are black squares and those bound to ≥50% exonic sequence are blue circles. ASOs bound to 5 exonic regions induce skipping >50% of the positive control ASO. The convention used for ASO naming has the locations of the 5’ and 3’ bound nts relative to the 5’ most nt of Exon 44. All 47 ASOs were tested at 4 μM with 6 replicates on plate replicate and then only the top performers were retested twice at 4 μM with 4 replicates per plate. The error bars represent the variability across three biological replicates.

**Figure 2.**
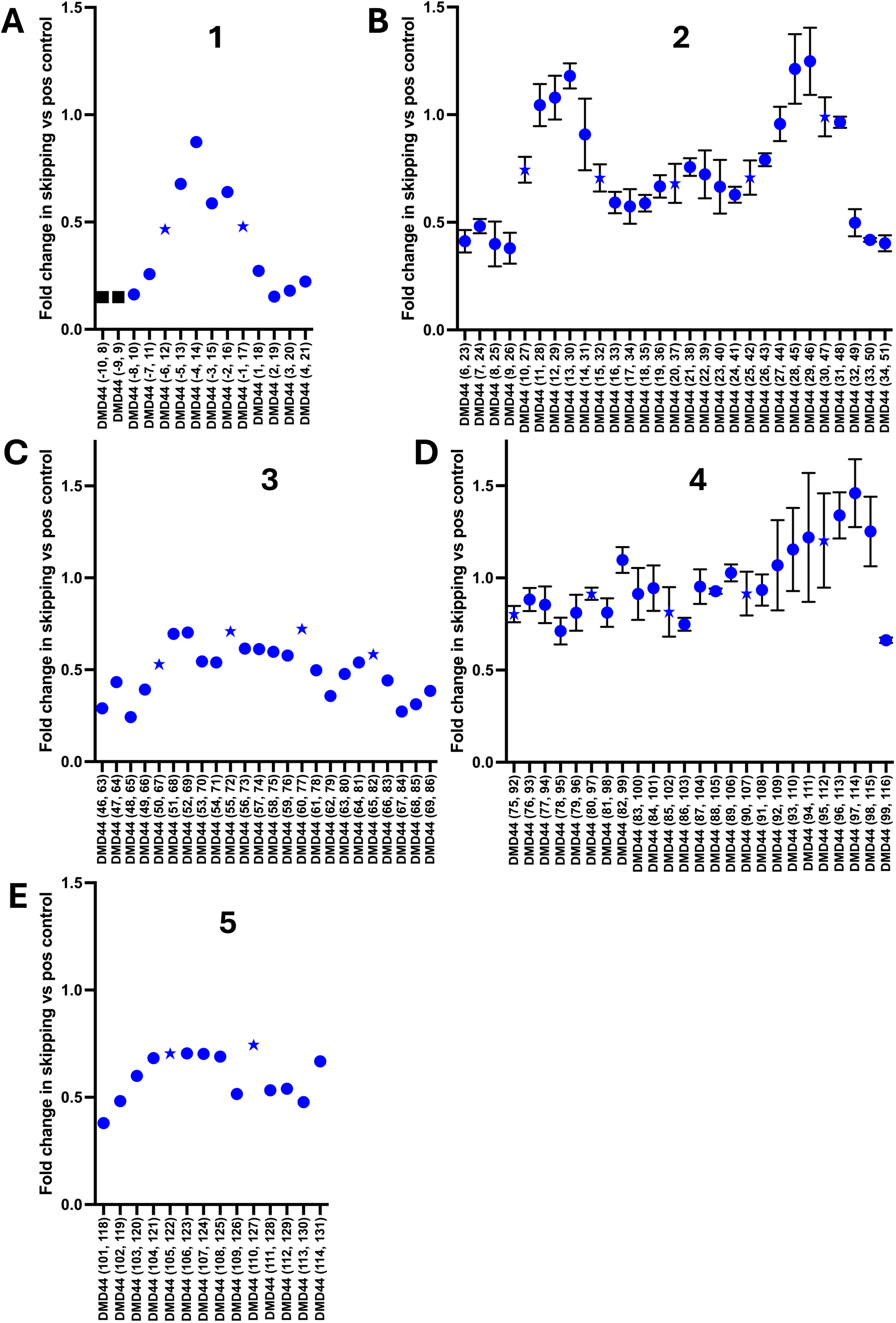
Microtiling of exon 44 refines optimal ASO target sites across five regions. Additional 18mer ASOs were designed to broaden coverage within regions 1–5 and tested in a gymnotic assay in Del42,43 myotubes. In region #1, DMD44 (−4, 14) showed ~2-fold higher activity than neighboring ASOs found in the primary screen (A). Region #2 microtiling revealed two distinct activity peaks and identified multiple ASOs with improved potency compared to the primary screen (B). Region #3 yielded no ASOs with improved activity but several that matched activity for the primary screen hits (C). In region #4, several ASOs were equipotent to the primary hits but none were clearly superior (D). Region #5 yielded no ASOs with improved activity, although several were equipotent to the primary screen hits. Overall, regions #2 and #4 showed the highest activity, followed by region #1, with regions #3 and #5 the least active. ASOs found in the primary screen are shown as stars. ASOs bound to ≥50% intronic sequence are black squares and those bound to ≥50% exonic sequence are blue circles.

### Validation of the primary course tiling and secondary microtiling screens

A set of 22 ASOs with a range of activities from the primary and secondary screens were selected to evaluate in two different cell lines to prevent the possibility that the results are specific to the Del42,43 cell line and thus less likely to translate to mouse models and eventually patients. The activities of these 22 ASOs were strongly correlated between the Del42,43 cell line and another DMD patient derived myoblast cell line (Del52, “KM571”) and an iPSC cardiomyocyte cell line (P = < 0.001, Fig. 3). This confirms that the Del42,43 cell line is suitable for screening for active exon 44 skipping ASOs and that the ASOs found therein have a high probability of working in the heart as well as skeletal muscle cells. As found in the Del42,43 cell line, regions #2 and #4 appear to be the most active in the Del52 myoblast and iPSC cardiomyocyte lines.

**Figure 3.**
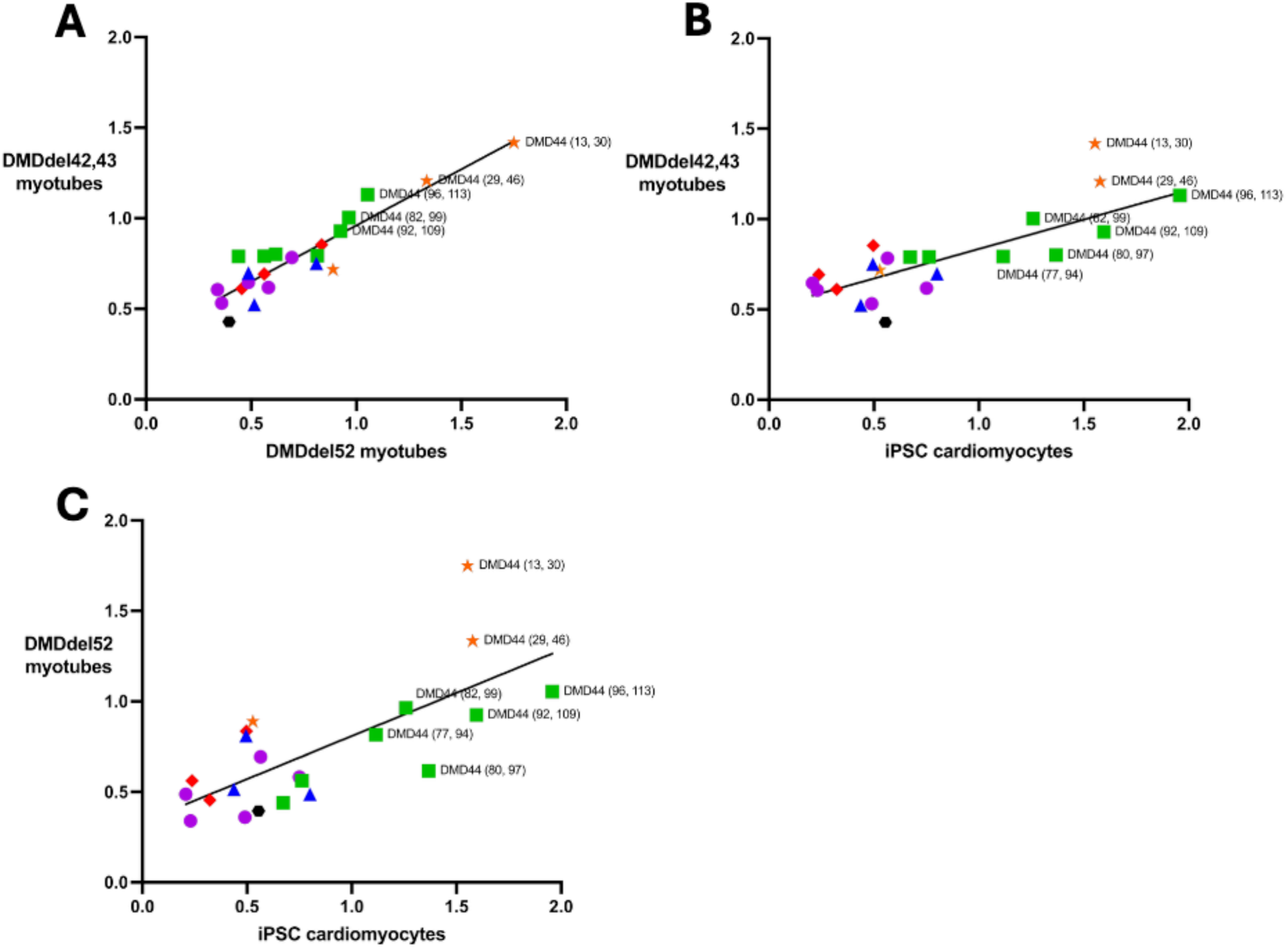
ASO activity is conserved across muscle and cardiac cell models. A set of 22 ASOs spanning a range of activities was evaluated in Del42,43 myotubes, a second DMD patient-derived myoblast line (Del52, KM571), and iPSC-derived cardiomyocytes. ASO activity was strongly correlated across cell types (P < 0.001), including 7796 vs KM571 (A), 7796 vs iPSC cardiomyocytes (B), and KM571 vs iPSC cardiomyocytes (C). Regions 2 and 4 showed the highest activity across all models.

### Analogs of top Exon 44 skipping ASOs with different lengths rarely had improved activity

Next, attempts were made to improve the activity of top ASOs targeting regions #2 and #4 by altering their lengths. Shorter ASOs (< 18 nts) might distribute into cells better leading to better potency, while longer ASOs (> 18 nts) might have better potency through stronger binding affinity to the target site. Families of shorter and longer versions of DMD44 (13, 30), DMD44 (29, 46), DMD44 (82, 99), DMD44 (96, 113), which are among the top ASOs targeting regions #2 and #4, were designed and tested in the Del42,43 myotubes via gymnosis.

There was a clear trend for the DMD44 (13, 30) family where 14mer and 15mer versions showed less activity than the parental 18mer DMD44 (13, 30) (Fig. 4A). Two of three 16mers tested, both 17mers, both 19mers, and two of three 20mers were equipotent to DMD44 (13, 30). In contrast, longer ASOs (21mers, 22mer, and 24mers) showed reduced activity compared to DMD44 (13, 30) (Fig. 4A). This suggests that the optimal ASO length for gymnotic assays range from 16 to 20 nts in length, though this might depend on cell line and ASO chemistry used.

**Figure 4.**
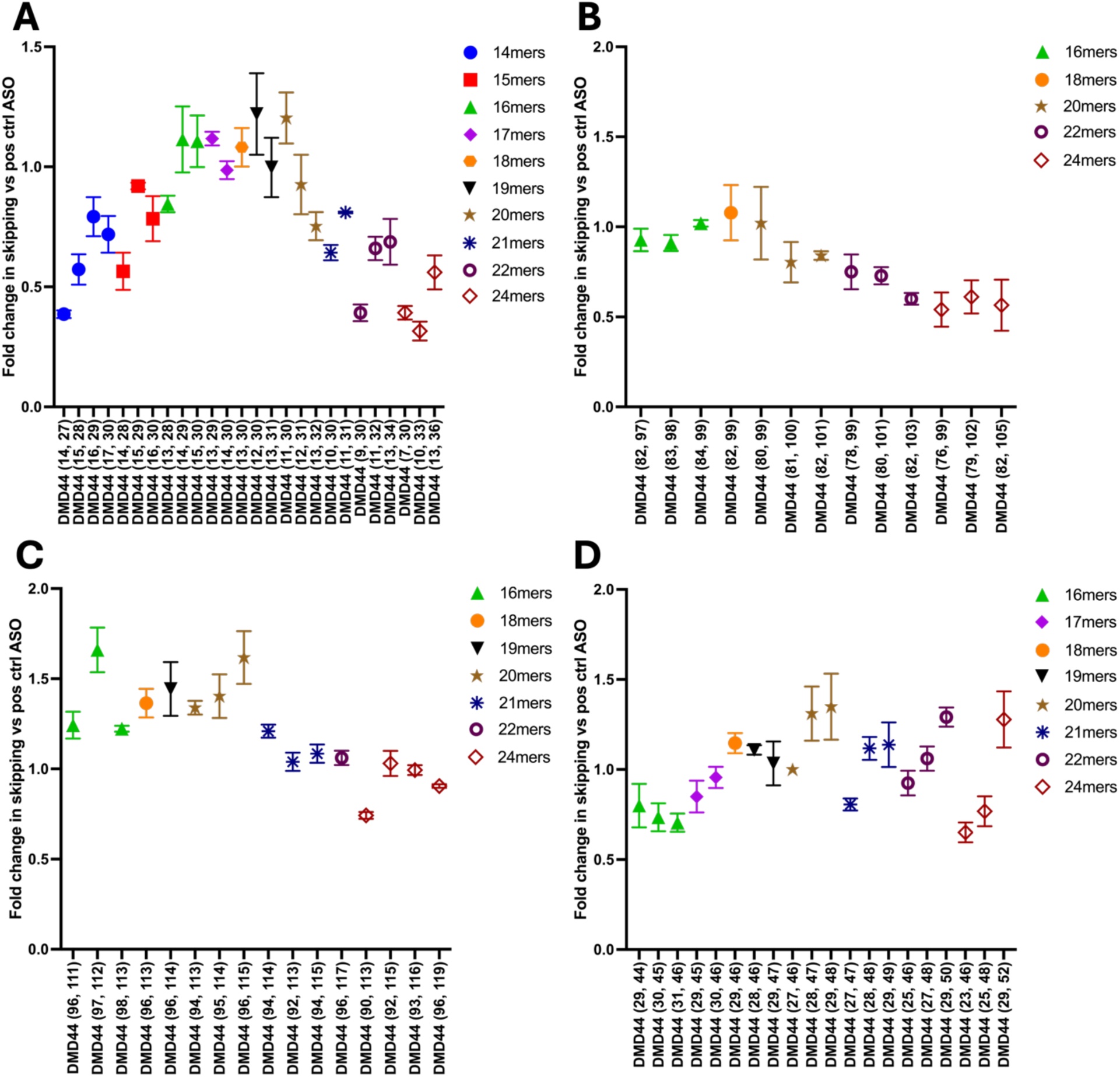
ASO length optimization identifies a narrow window for maximal exon 44 skipping activity. Length variants (14–24mers) of top ASOs targeting regions #2 and #4 were evaluated in Del42,43 myotubes. For DMD44 (13, 30), shorter ASOs (≤15mer) showed reduced activity, while 16–20mers were generally equipotent and longer ASOs (≥21mer) were less active (A). Similar trends were observed for DMD44 (82, 99) and DMD44 (96, 113) (B, C). In contrast, the DMD44 (29, 46) family showed reduced tolerance to shorter ASOs, but greater tolerance to longer analogs (D). Across 72 analogs tested, only one shorter ASO (16mer DMD44 (97, 112)) demonstrated improved activity over its 18mer parent. These data support an optimal PS/2’MOE ASO length of approximately 16–20 nts for exon 44 skipping.

The DMD44 (82, 99) and DMD44 (96, 113) families (Figs. 4B and 4C) were like the DMD44 (13, 30 family) (Fig. 4A). For these two families, 16mers, 19mers, and 20mers were similarly potent to the 18mer parent ASO but 21mers, 22mers, and 24mers were less potent.

The DMD44 (29, 46) family was unique. For this family, shorter analogs were less tolerated while longer analogs were more tolerated. The 16mers and 17mers were less potent than DMD44 (29, 46) and the 19mers, 2 of 3 20mers 2 of 3 21mers, and 1 of 3 24mers were similarly potent to DMD (29, 46) (Fig. 4D).

Of the 72 analogs tested with lengths ranging from 14 to 24 nts, only 1 ASO appeared to have clearly better potency than the 18mer parent ASO, the 16mer DMD44 (97, 112) (Fig. 4C). It may be that shorter ASOs rarely work better and are often worse than 18mers because shorter ASOs have reduced binding affinity and may be gaining more off target binding opportunities, which would reduce the overall amount of ASO available to bind exon 44. Longer ASOs may rarely be better than 18mers because more binding affinity may provide negligible benefit as the affinity of an 18mer to target is already high. Alternatively, ASOs longer than 20 nts may have decreased activity because their higher binding affinity may reduce availability of an ASO to bind and skip other transcripts’ exon 44, longer ASOs may have a higher probability of self-dimerization, more potential for protein binding due to a longer PS backbone and could have reduced productive uptake through diminished cellular uptake, reduced endosomal escape, or poorer ability to navigate to the nucleus.

### Nucleofection restored activity of longer ASOs that worked less well via gymnosis

A trend that was observed during ASO optimization was that ASOs longer than 20 nts in length often had diminished activity compared to analogs which were of 16-20 nts in length. One possible reason for the decreased activity could be that longer ASOs have more difficulty with productive uptake through either hampered endocytosis, reduced endosome escape, or diminished navigation to the nucleus. To test this, a set of ASOs from the DMD44 (13, 30) and DMD44 (29, 46) families were tested via nucleofection, an electroporation method that delivers ASOs to the nucleus. For the DMD44 (13,30) family (Fig. 5A), nucleofection improved the activity of both 24mers tested compared to what was observed via gymnosis. The same occurred for one of the 22mers (DMD44 (9, 30)) and a 21mer (DMD44 (10,30)). This implies that for these 4 ASOs, a reason for their drop-in activity in the gymnotic assay compared to shorter analogs is due to their larger size having a negative impact on productive uptake (Fig. 5A). The DMD44 (29, 46) family behaved similarly where the activities of one 20mer, 22mer, and 24mer were increased when tested via nucleofection compared to what was observed by gymnosis implying that they too had hindered productive uptake due to their greater length (Fig. 5B).

**Figure 5.**
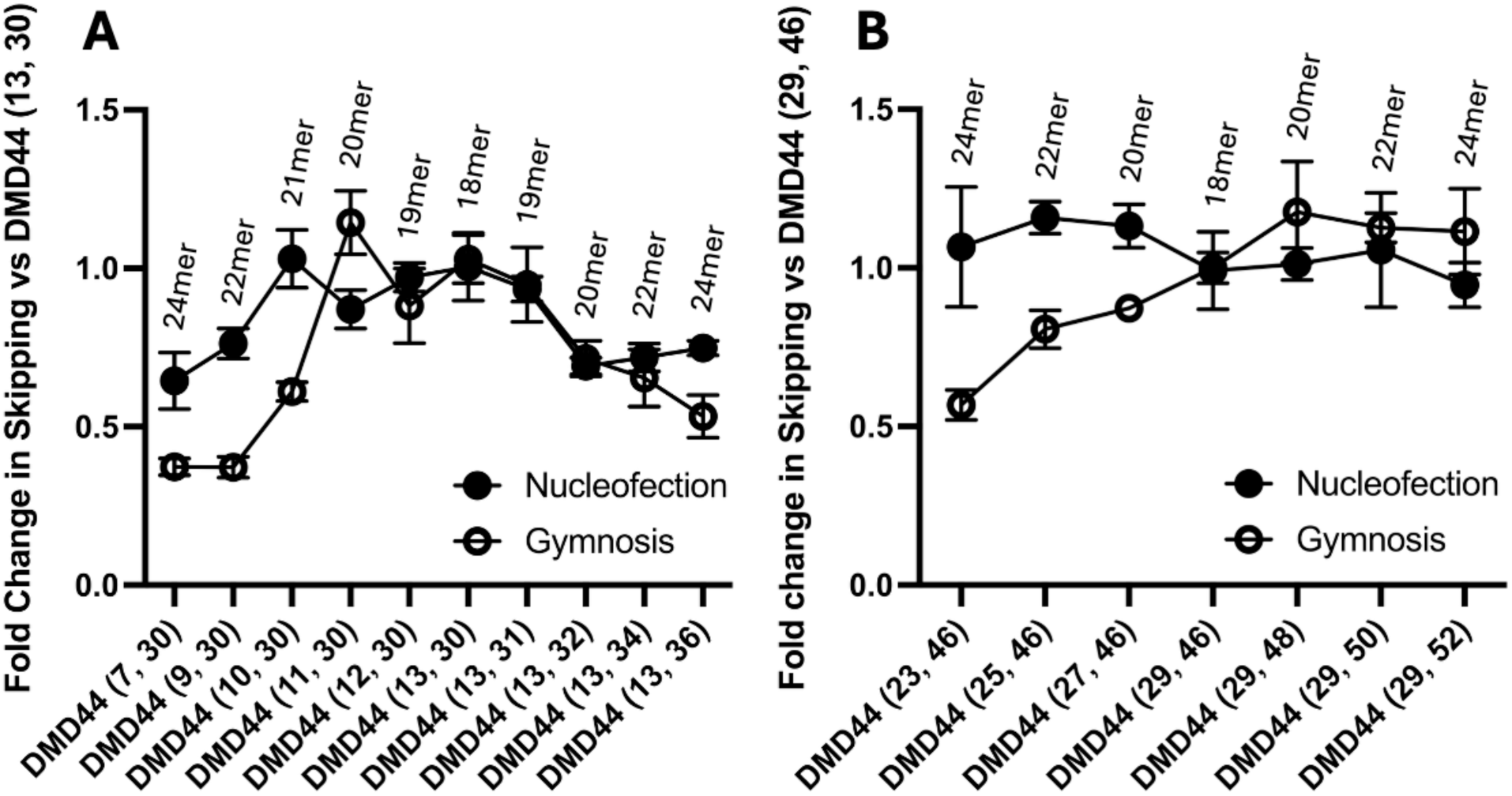
Nucleofection restores activity of longer ASOs with reduced gymnotic uptake. ASOs from the DMD44 (13, 30) and DMD44 (29, 46) families were evaluated by nucleofection to assess whether reduced activity of longer ASOs (>20mers) in gymnotic assays was due to impaired productive delivery. In the DMD44 (13, 30) family, nucleofection increased activity of multiple 21-24mer ASOs that were less active under gymnotic conditions (A). In the DMD44 (29, 46) family, activity of multiple 20-24mer ASOs was similarly restored by nucleofection (B). Across both families, lengthening ASOs did not increase activity beyond the parental 18mers, indicating that additional length provides minimal benefit to on-target potency.

Nucleofection did not recover activity for all ASOs that had lower activities compared to the parental 18mer in the gymnotic assay (Fig. 4). The 20mer DMD44 (13,32) and the 22mer DMD44 (13, 34) did not reach the activity of the parental 18mer when tested by nucleofection. It may be that these ASOs have some other property, such as secondary structures or self-self structures that reduces on-target activity, or they may cover other regulatory sites on exon 44 leading to greater exon inclusion. In addition, none of the longer analogs surpassed the activity of the 18mer in the nucleofection assay. This may be because the 18mer PS/2’MOE design is suitable for full activity for these sequences and adding more nts provides only negligible benefit.

### The addition of terminal LNAs rarely improved activity and can also have negative consequences

To try and increase the activity of ASOs from the DMD44 (13, 30), DMD44 (29, 46), DMD44 (82, 99), DMD44 (96, 113) families single terminal locked nucleic acids (LNAs) were added to the 5’ and 3’ positions (Fig. 6). LNAs increase binding affinity and terminal LNAs can improve ASO stability by blocking exonuclease activity [27]. For the 17 ASOs designed to have terminal LNAs, only 1 showed improved activity over the non-LNA version (DMD44 (30, 47) LNA), (Fig. 6D). Reciprocally, there were two instances where terminal LNAs led to a significant drop in activity compared to the non-LNA versions (DMD44 (13, 30) LNA and DMD44 (97, 112) LNA (Figs. 6A and 6C). For the other 14 ASOs with terminal LNAs, there was no apparent impact on activity compared to the LNA-free forms. This suggests that fully 2’ MOE PS ASOs have suitable binding affinity to induce exon skipping and increasing affinity with the addition of LNAs rarely has a positive impact and can just as frequently lead to a negative impact.

**Figure 6.**
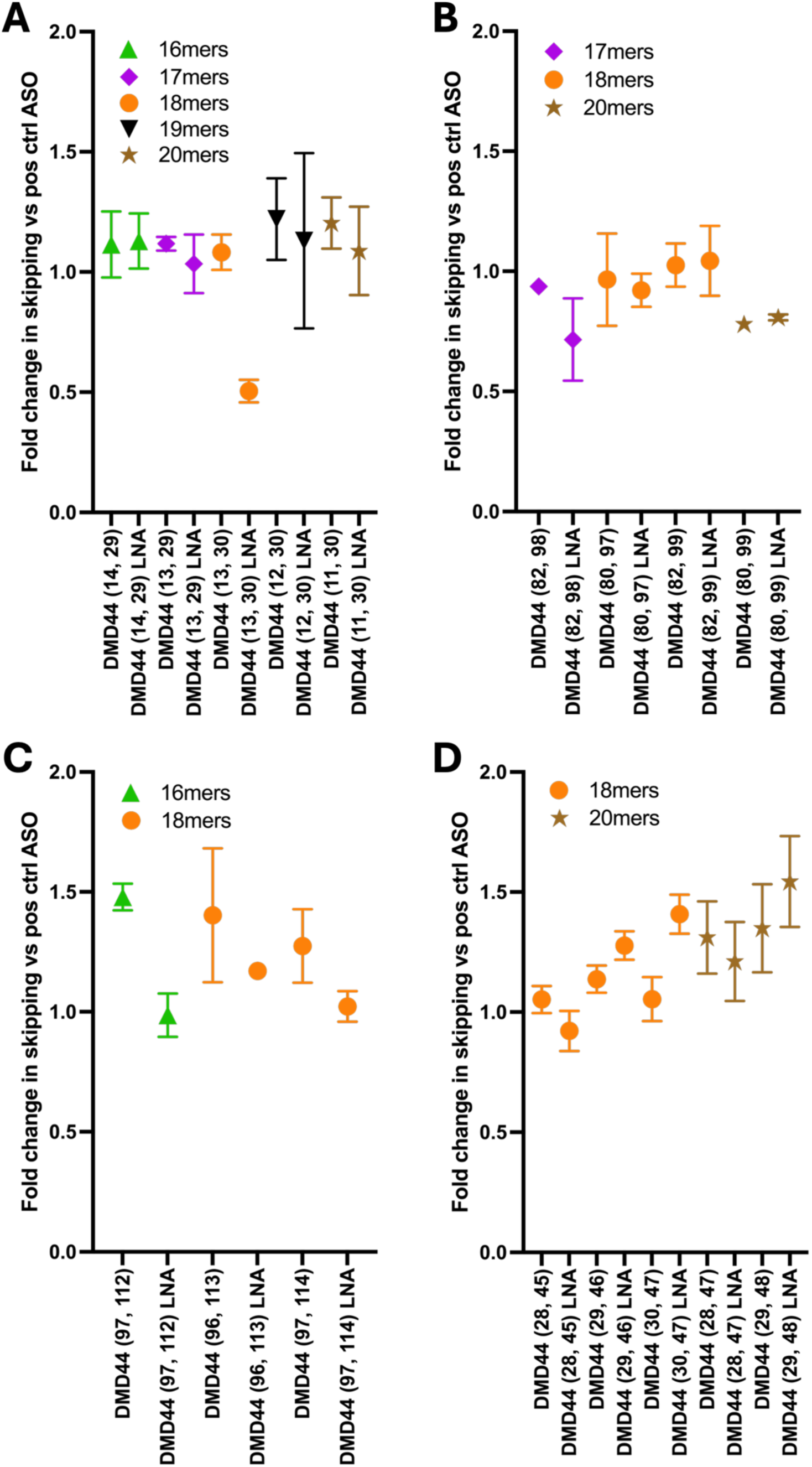
Terminal LNA modifications have limited impact on ASO activity. Single locked nucleic acids (LNAs) were added to the 5′ and 3′ termini of ASOs from the DMD44 (13, 30) (A), DMD44 (82, 99) (B), DMD44 (96, 113) (C), and DMD44 (29, 46) (D) families to assess effects on exon 44 skipping. Across 17 LNA-modified ASOs, only one (DMD44 (30, 47) LNA) showed improved activity relative to the parent ASO (B). Two ASOs exhibited reduced activity with terminal LNA incorporation (DMD44 (13, 30) LNA and DMD44 (97, 112) LNA) (A, C), while the remaining ASOs showed no consistent change. These data indicate that terminal LNA addition rarely enhances activity and can negatively impact exon skipping, suggesting that fully 2′-MOE PS 18mer ASOs have sufficient binding affinity for activity.

### DMD44 (82, GG) LNA was found to be the top performing ASO *in vivo* among a set among 14 tested

To determine which ASOs from the DMD44 (13, 30), DMD44 (29, 46), DMD44 (82, 99) and DMD44 (96, 113) families have the most potential in vivo, top performing ASOs from each family were tested in a mouse model carrying 4 copies of the human *DMD* gene integrated in mouse chromosome 5 [20]. In the first study (Figs. 7A–7D) ASOs were dosed IV via tail vein at 1.71 µmol/kg. Animals were dosed weekly, for 4 weeks, with dosing days on day 0, day 7, day 14, day 21. Animals were sacrificed on day 28 and the quadricep and heart tissues were harvested. This short dosing regimen was utilized as a rapid way to test and evaluate ASOs *in vivo*. Groups of animals were dosed with either vehicle or an exon 51 skipping ASO (AON-C12) as negative controls. In the second study, the same weekly dosing for 4 weeks regimen was applied, but the dose of the ASOs was increased to 3.42 µmol/kg (Figs. 7E and 7F).

**Figure 7.**
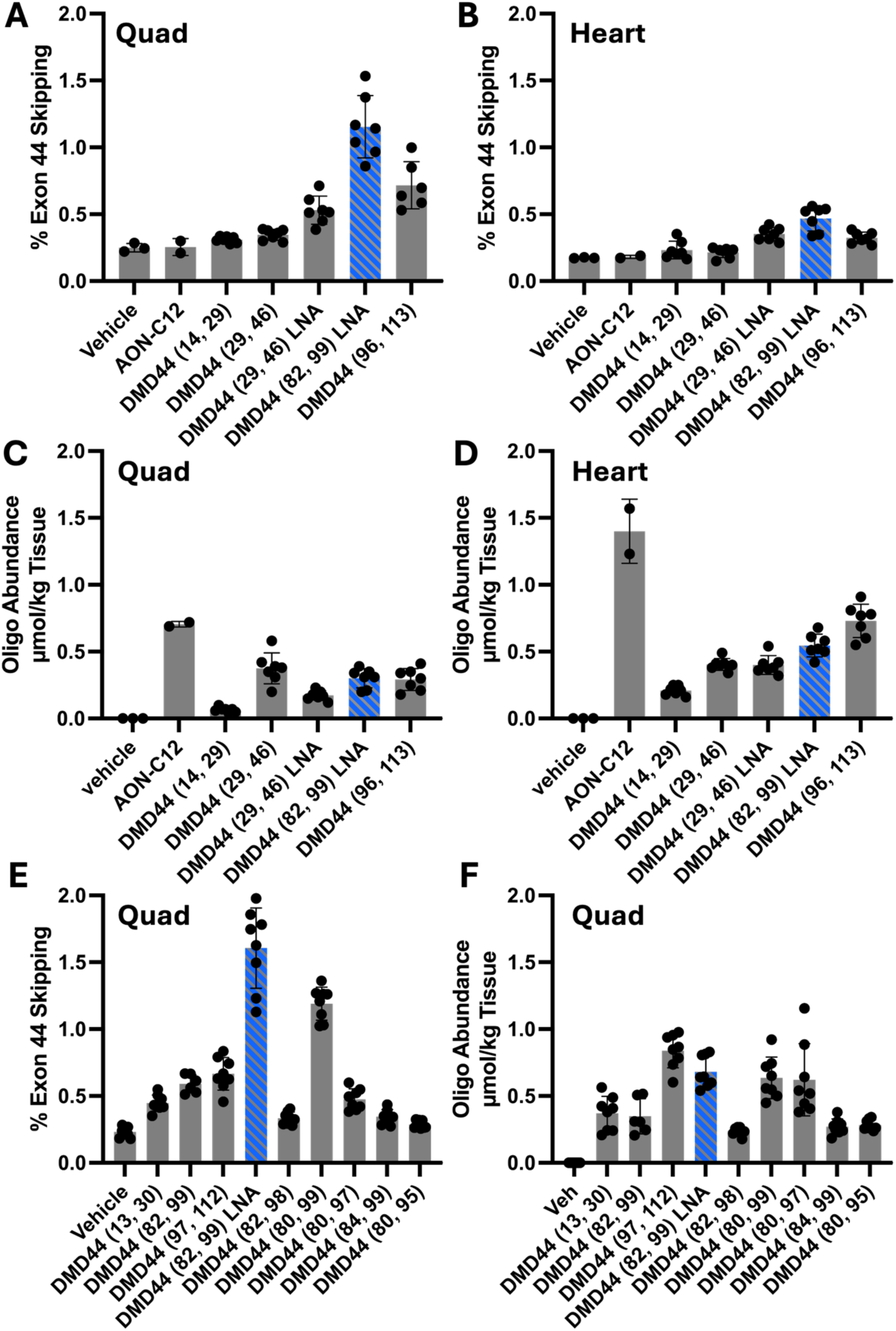
In vivo evaluation identifies DMD44 (82, GG) LNA as the top-performing exon 44 skipping ASO. Top ASOs from the DMD44 (13, 30), DMD44 (29, 46), DMD44 (82, 99), and DMD44 (96, 113) families were evaluated in a humanized DMD mouse model following weekly intravenous dosing for 4 weeks. In study 1 (1.71 µmol/kg), exon 44 skipping and tissue levels were measured in quadriceps (quad) (A, C) and heart (B, D). DMD44 (82, 99) LNA showed the highest activity (~1.2% in quadriceps and ~0.4% in heart), while the exon 51 control ASO (AON-C12) did not induce exon 44 skipping. In study 2 (3.42 µmol/kg), ASOs were evaluated in quadriceps (E, F), and DMD44 (82, 99) LNA remained the top performer, showing the most exon skipping. Consistent with the higher dose compared to study 1, its tissue levels were elevated in study 2 and it also outperformed its full 2’MOE counterpart.

In study 1, 5 exon 44 ASOs from the DMD44 (13, 30), DMD44 (29, 46), DMD44 (82, 99), DMD44 (96, 113) families were tested and DMD44 (82, 99) LNA emerged as the best showing ~1.2% skipping in the quadriceps and ~0.4% skipping in the heart. This low level of skipping was expected given the short duration of the study, the fact that exon 44 skipping creates and out of frame transcript in this model and that ASOs deliver less to healthy muscle tissue. Exon 51 skipping AON-C12 did not induce exon 44 skipping over levels observed in the vehicle group. Tissue levels of the ASOs were measured via LCMS. DMD44 (14, 29) was an outlier with the lowest quadriceps and heart levels. The negative control ASO AON-C12 had the highest tissue levels, though notably its chemistry is slightly different. This fully PS 18mer has 14 2’ OMe and 4 LNA modifications. Perhaps the altered chemistry enabled its higher tissue levels. The levels of non-skipped dystrophin mRNA were not impacted by the treatments relative to vehicle (Fig. S1).

In study 2, DMD44 (82, 99) LNA was tested again along with 8 other top performers across the 4 families at double the concentration of what was tested in study 1. DMD44 (82, 99) LNA was again the top performing ASO. Consistent with doubling the dose, the exon 44 skipping and tissue levels of DMD44 (82, 99) LNA doubled compared to what was observed in study 1. DMD44 (82, 99) LNA also outperformed its LNA-free version (DMD44 (82, 99)) in exon skipping and tissue levels. The levels of non-skipped dystrophin mRNA were not changed from the treatments relative to vehicle (Fig. S2).

### Activity of DMD44 (82, GG) LNA versus PMO comparator in a DMDdel45/*mdx* mouse model

To further study the potential of the best performing DMD44 (82, 99) LNA, it was next tested in the DMDdel45/*mdx* mouse model. This mouse carries a deletion of exon 45 in all its copies of the *hDMD* gene, in a mouse dystrophin-deficient background (*mdx*). A deletion of exon 45 is the most common exon 44 skip amenable mutation in patients. Due to the lack of human and mouse dystrophin this mouse model also has a dystrophic phenotype making it suitable to study exon 44 skipping ASOs in a disease relevant background and allowing better ASO delivery to muscle. A comparator exon 44 skipping PMO ASO was also evaluated in this study [15]. ASOs were dosed intravenously via the tail vein weekly, for 4 weeks, with dosing days on day 0, day 7, day 14, day 21. Animals were sacrificed on day 28 and quadriceps muscles were harvested on day 28. This matches the dosing paradigm applied in the two studies in the hDMD mouse (Fig. 7). DMD44 (82, 99) LNA was dosed at 3.18 µmol/kg and 0.636 µmol/kg. The comparator PMO was dosed at 3.18 µmol/kg and at a 5-fold higher dose (15.9 µmol/kg) to reflect clinically relevant dosing, where PMOs are typically administered at higher levels than PS ASOs due to their reduced bioavailability.

Exon 44 skipping was assessed by RT-PCR (Fig. 8A) and ddPCR (Fig. 8B). Trends were consistent across both methods, although RT-PCR values were ~2–4-fold higher than ddPCR, consistent with known overestimation by RT-PCR, which remains suitable for rank ordering ASO potency but not for absolute quantification of exon skipping levels. Both ASOs showed dose-dependent exon skipping. The comparator ASO at 15.9 µmol/kg induced the highest level of skipping (22.8% by ddPCR; Fig. 8B). At 3.18 µmol/kg, the comparator ASO showed slightly higher activity (15.3% by ddPCR) compared to DMD44 (82, 99) LNA (14.4%; Fig. 8B), while DMD44 (82, 99) LNA at 0.636 µmol/kg induced the lowest level of skipping (4.6%; Fig. 8B).

**Figure 8.**
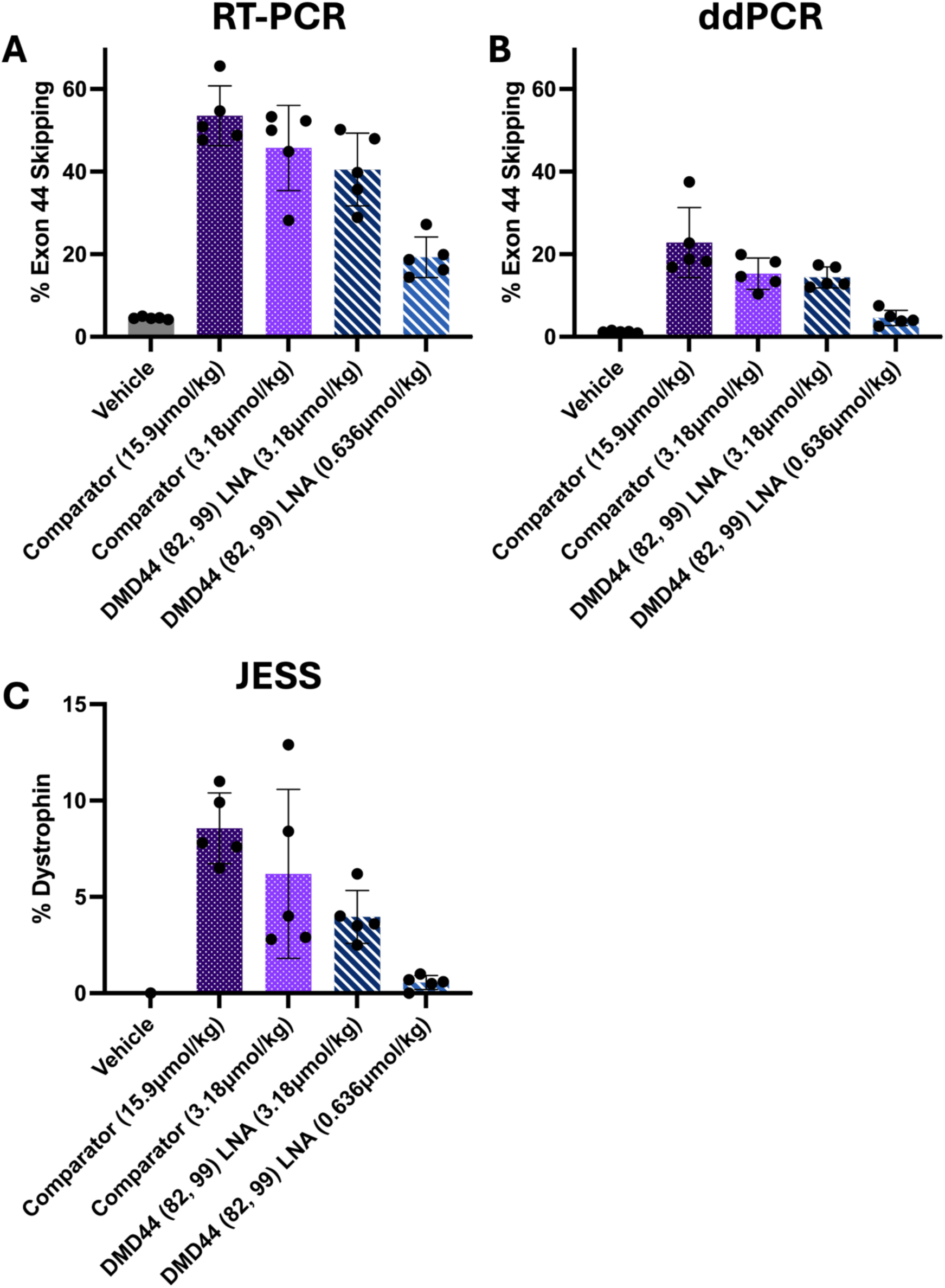
In vivo activity of DMD44 (82, GG) LNA compared to an exon 44 skipping PMO in the DMDdel45/mdx mouse model. DMD44 (82, 99) LNA and a comparator PMO ASO were evaluated in DMDdel45/mdx mice following weekly IV dosing for 4 weeks. Exon 44 skipping in quadriceps was assessed by RT-PCR (A) and ddPCR (B), and dystrophin protein levels were quantified relative to normal controls (C). Both ASOs showed dose-dependent exon skipping and dystrophin restoration. The comparator PMO at 15.9 µmol/kg induced the highest levels of exon skipping (~22.8% by ddPCR) and dystrophin (~8.6%), while at 3.18 µmol/kg both ASOs showed comparable activity. DMD44 (82, 99) LNA at 0.636 µmol/kg showed reduced activity. RT-PCR values were consistently higher than ddPCR but showed similar trends across groups.

Dystrophin restoration was also determined by comparing the amounts of dystrophin protein in the treatment groups to a standard curve (Fig. 8C). The trends in dystrophin protein production matched those seen for exon skipping (Figs. 8A, 8B). The comparator ASO at the high dose (15.9 µmol/kg) induced the highest level of dystrophin protein at 8.6% (Fig. 8C). At 3.18 µmol/kg, the comparator ASO trended to be slightly better with 6.2% compared to DMD44 (82, 99) LNA which induced 4.0% (Fig. 8C). DMD44 (82, 99) LNA at 0.636 µmol/kg induced the lowest amount of dystrophin production at 0.6% (Fig. 8C).

At face value, these data suggest that the comparator PMO may be more active than DMD44 (82, 99) LNA, based on higher levels of exon skipping and dystrophin production. This difference was most evident at the 5-fold higher dose (15.9 µmol/kg) compared to 3.18 µmol/kg DMD44 (82, 99) LNA, a dose level difference that is clinically relevant given the lower bioavailability but also greater tolerability of PMOs. However, direct comparison between PMOs and PS ASOs *in vivo* is not straightforward, as PS ASOs tend to accumulate in tissues to a greater extent upon repeat dosing. As a result, the activity observed after 4 weeks is likely to underestimate the longer-term potential of PS ASOs relative to PMOs [28, 29].

## Discussion

### *In vitro* cellular screening identified active regions of exon 44 and the best 18mer PS/2’MOE ASOs that target them

To identify the top regions to target to induce DMD exon 44 skipping, a primary screen was carried out with forty-seven 18mer PS/2’MOE ASOs that step 5 nts at a time starting 50 nts upstream of exon 44 in intron 43 and ending 49 nts downstream of exon 44 in intron 44 (Fig. 1). This primary screen was carried out via gymnotic uptake in Del42,43 patient derived myotubes, a very physiologically relevant system. Of the 47 ASOs, 83% induced exon 44 skipping. There were 5 regions that carried the most active ASOs. Region #1 spans positions −6 to 17 (23 nts), region #2 spans positions 10-47 (38 nts), region #3 spans positions 50-82 (33 nts), region #4 spans positions 75-112 (38 nts), and region #5 spans 105-127 (23 nts). These regions must function to promote exon 44 inclusion. When ASOs are bound to them, they may disrupt mRNA secondary structure which promotes exon inclusion. Alternatively, these regions may carry exonic splice enhancers (ESEs) that are bound by RNA binding proteins, such as the serine arginine rich (SR) proteins, which function to promote exon inclusion [30]. ASOs bound to these regions may block SR or other protein binding to ESEs and induce exon skipping.

A secondary screen was then carried out where each region was microtiled with 18mer PS/2’MOE ASOs that stepped across them 1 base at a time (Fig. 2). Region #1 saw the biggest improvement with DMD44 (−4, 14) being ~2X better than neighboring top ASOs from the primary screen. Interestingly, region #2 split into two regions after microtiling, with 5 ASOs emerging as better than neighboring top ASOs from the primary screen. The secondary screen revealed several ASOs from regions #3, #4, and #5 that were equipotent to those from the primary screen, but none that were clearly better.

Region #4 appeared to be the best to target to induce Exon 44 skipping followed by region #2, region #1, and lastly regions #3 and #5, which were similar. Presumably regions #4 and #2 play the biggest role in exon 44 inclusion. That trend was confirmed when a set of ASOs were tested in two different cell lines; another DMD patient derived myotube model (Del52) and an iPSC cardiomyocyte line (Fig. 3). In those cells, targeting regions #4 and #2 induced the highest amount of exon 44 skipping. In general, the potency of ASOs from across the 3 cell lines was strikingly similar, suggesting that ASO skipping potency, and exon 44 splicing regulation, is very similar from cell line to cell line.

Two key learnings emerged from this process. First, inclusion of an on-plate positive control ASO was critical for the screening workflow. Normalizing ASO activity to the positive control reduced plate-to-plate variability across experiments conducted over several weeks and is how data are presented in Figs. 1–4 and 6. This approach likely helped control for variability in gymnotic uptake arising from subtle differences in cell density or differentiation state, which are difficult to keep fully consistent. The second learning is that the 5 nt walk was successful in revealing the regions of top activity across the exon, with one caveat – the right side of region #2 may not have been found if the 5 nts walk had started 2 nts downstream from where the primary screen started. There is a steep drop in activity from DMD44 (30, 47) to DMD44 (32, 49). Therefore, for targeting regions such as exon 44, which is 148 nts in length, it may be most prudent to single base walk the entire exon if resources permit. Alternatively, a smaller step size such as a 3 bp walk with 18mers may be a good compromise.

### *In vitro* optimization revealed optimal length of PS/2’MOE ranging from 16 to 20bps and addition of terminal LNAs rarely provided benefit

The primary and secondary screens were performed using 18mer PS/2′MOE ASOs, consistent with the chemistry and length of Nusinersen, an FDA-approved splice-modulating ASO for SMA [24]. After identifying regions 4 and 2 as the most active, length variants of four top ASOs targeting these regions were designed and evaluated. These analogs ranged from 14 to 24 nts and maintained the PS/2′MOE chemistry. Of the 72 analogs tested, only one, the 16mer DMD44 (97, 112), showed improved activity relative to its 18mer parent.

A general trend was observed in which ASOs shorter than 16 nts and longer than 20 nts showed reduced activity (Fig. 4). Reduced activity of shorter ASOs may reflect decreased binding affinity to the target site or increased off-target binding, which could lower the effective concentration bound to exon 44. For longer ASOs, reduced activity may result from increased probability for secondary structure, diminished recycling due to higher binding affinity, or impaired gymnotic uptake, including reduced endocytosis or endosomal escape. Consistent with impaired gymnosis, the activity of several longer ASOs was restored by nucleofection, suggesting that reduced gymnotic uptake contributes to their lower potency (Fig. 5). This may be due to less efficient endocytic uptake.

To try and improve ASO activity, a set of top performing PS/2’MOE ASOs were modified by swapping the 2’MOE modification with an LNA modification at the 5’ and 3’ terminal positions (Fig. 6). For the 17 ASOs tested, only 1 ASO with terminal LNAs had a greater potency compared to its LNA-free parent. On the other hand, terminal LNAs reduced the activity of 2 ASOs compared to their LNA-free counterparts, and the other 14 ASOs with terminal LNAs had no change compared to their LNA-free counterparts. LNAs have a higher binding affinity compared to 2’MOE and may also have a higher resistance to exonuclease activity. Perhaps the additional binding affinity or stability provided by the terminal LNAs provides only negligible benefits over a fully PS/2’MOE design, which already has a high binding affinity and stability. Terminal LNAs could also reduce activity of ASOs by increasing the probability of intermolecular structures that reduce the effective concentration of ASO for exon 44 target site binding.

These optimization studies suggest that, for PS/2′MOE exon-skipping ASOs, lengths of 16 - 20 nts are generally optimal for potency, making 18mers a logical starting point for screening. In practice, it was observed that improving activity by altering length or adding terminal LNAs was rare. As a result, it may be most efficient to prioritize screening of 18mer PS/2′MOE ASOs and defer optimization until lead sequences demonstrate *in vivo* potential. That said, when resources permit, parallel *in vitro* optimization may accelerate lead identification by ensuring that the best analogs are already defined once sequences with in vivo potential are identified. Importantly, identifying a set of analogs targeting the same site through *in vitro* studies remains valuable, as subtle design differences can translate to improved *in vivo* performance, as observed for DMD44 (82, 99) LNA (Fig. 7), which outperformed its LNA-free counterpart despite similar *in vitro* activity.

### All top ASOs found from *in vitro* studies induced Exon 44 skipping *in vivo* and DMD44 (82, GG) LNA emerged as the top performer with its potential in DMD revealed through comparison to a relevant comparator PMO

Top performing ASOs from the region #2 targeting DMD44 (13, 30) and DMD44 (29, 46) families and the region #4 targeting DMD44 (82, 99) and DMD44 (96, 113) families were tested in a mouse model carrying 4 copies of a full human DMD transgene. A 4-week study paradigm was established as a quick approach to rank order ASOs for *in vivo* potency. It’s possible that shorter regimens would also be sufficient. The dose levels of 1.71µmol/kg and 3.42 µmol/kg range roughly from 10 to 30 mg/kg, which are doses of PS ASOs known to typically be well tolerated in mice.

All of the 14 ASOs induced exon 44 skipping, with DMD44 (82, 99) LNA being the top performer showing 1.2% skipping in quad (Fig. 6A) and 0.45% skipping in heart (Fig. 6B) when dosed at 1.71 µmol/kg. At 3.42 µmol/kg, DMD44 (82, 99) LNA induced 1.6% skipping in the quad, which when taking into account % skipping over background levels, is about a 2-fold increase in activity compared to the 1.71 µmol/kg dose indicating a good correlation between dose level and activity (Fig. 6B). Tissue levels were also dose-responsive going from ~0.5 µmol/g in the quad of the 1.71 µmol/kg dosed animals (Fig. 6D) to ~0.75 µmol/kg in the quad of the 3.42 µmol/kg dosed animals. Interestingly, although DMD44 (82, 99) LNA and its LNA-free version (DMD44 (82, 99)) had indistinguishable *in vitro* potencies, DMD44 (82, 99) LNA was clearly superior *in vivo* (Fig. 6E). This may be because DMD44 (82, 99) LNA had fold higher tissue levels, which is a property that may not always be selected for *in vitro* assays. This suggests that once the best analogs are found from *in vitro* assays several should be tested *in vivo* even if their *in vitro* activities are the same. Surprising differences might be revealed once evaluated *in vivo*.

DMD44 (82, 99) LNA was also evaluated alongside a PMO comparator in the disease-relevant hDMDdel45/*mdx* mouse model (Fig. 8). Direct comparison between PS ASOs and PMOs *in vivo* is not straightforward, as PS ASOs accumulate in tissues with repeat dosing, whereas PMOs do not, making short-term studies more likely to underestimate PS ASO activity [28, 29]. In contrast, PMOs can be administered at higher doses due to their improved tolerability. To partially account for these differences, PMOs were dosed at up to 5-fold higher levels than PS ASOs, with one shared dose level. At the high dose (15.9 µmol/kg), the PMO induced the highest exon skipping (22.8% by ddPCR; Fig. 8B) and dystrophin production (8.6%; Fig. 8C), while DMD44 (82, 99) LNA induced 14.4% skipping and 4.0% dystrophin at 3.18 µmol/kg. These dose levels are relevant for comparison given clinical dosing practices, although the activity of DMD44 (82, 99) LNA may be underestimated in this short-term study due to its expected accumulation with repeat dosing. Longer-term studies would be needed to more fully assess its relative potential.

Exon 44 skipping was evaluated using both RT-PCR (Fig. 8A) and ddPCR (Fig. 8B). The results showed a consistent trend across all groups, but the percentage of skipping measured by RT-PCR was roughly two to four times higher than what as observed with ddPCR. It’s well established that RT-PCR tends to overestimate skipping percentages, though it remains suitable for rank ordering the potency of ASOs.

## Conclusion

This work identified multiple active regions within DMD exon 44 that can be targeted to induce robust exon skipping and showed that the relative activity of ASOs is highly consistent across skeletal muscle and cardiac cell models. Our *in vitro* optimization studies further suggest that, for fully PS/2′MOE steric-blocking ASOs, lengths of 16 - 20 nts are generally preferred, whereas longer designs and terminal LNA incorporation rarely improve activity and can sometimes be detrimental. *In vivo* testing showed that all top ASOs retained activity, with DMD44 (82, 99) LNA emerging as the strongest performer across the mouse studies we conducted. Comparison to a relevant PMO comparator in the hDMDdel45/*mdx* model also provided useful context for interpreting the activity of this chemistry in a disease-relevant setting. Together, these findings define practical design principles for exon 44 targeted ASOs and highlight the importance of evaluating both sequence and chemistry early in discovery. More broadly, the approach described here may help guide development of future exon 44 skipping programs and could also be informative for other steric-blocking ASO efforts.

## Supplementary information

**Table S1.**
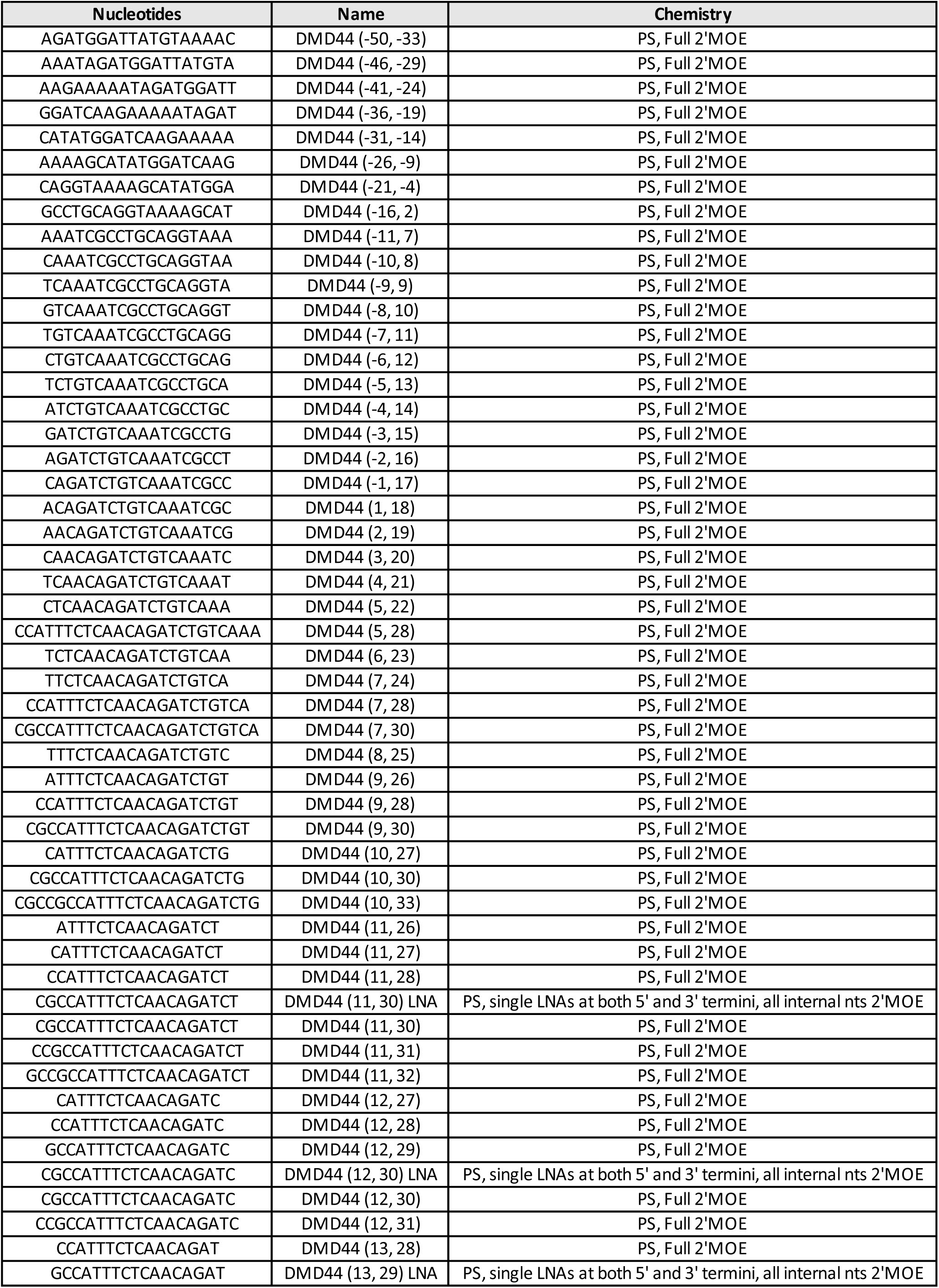
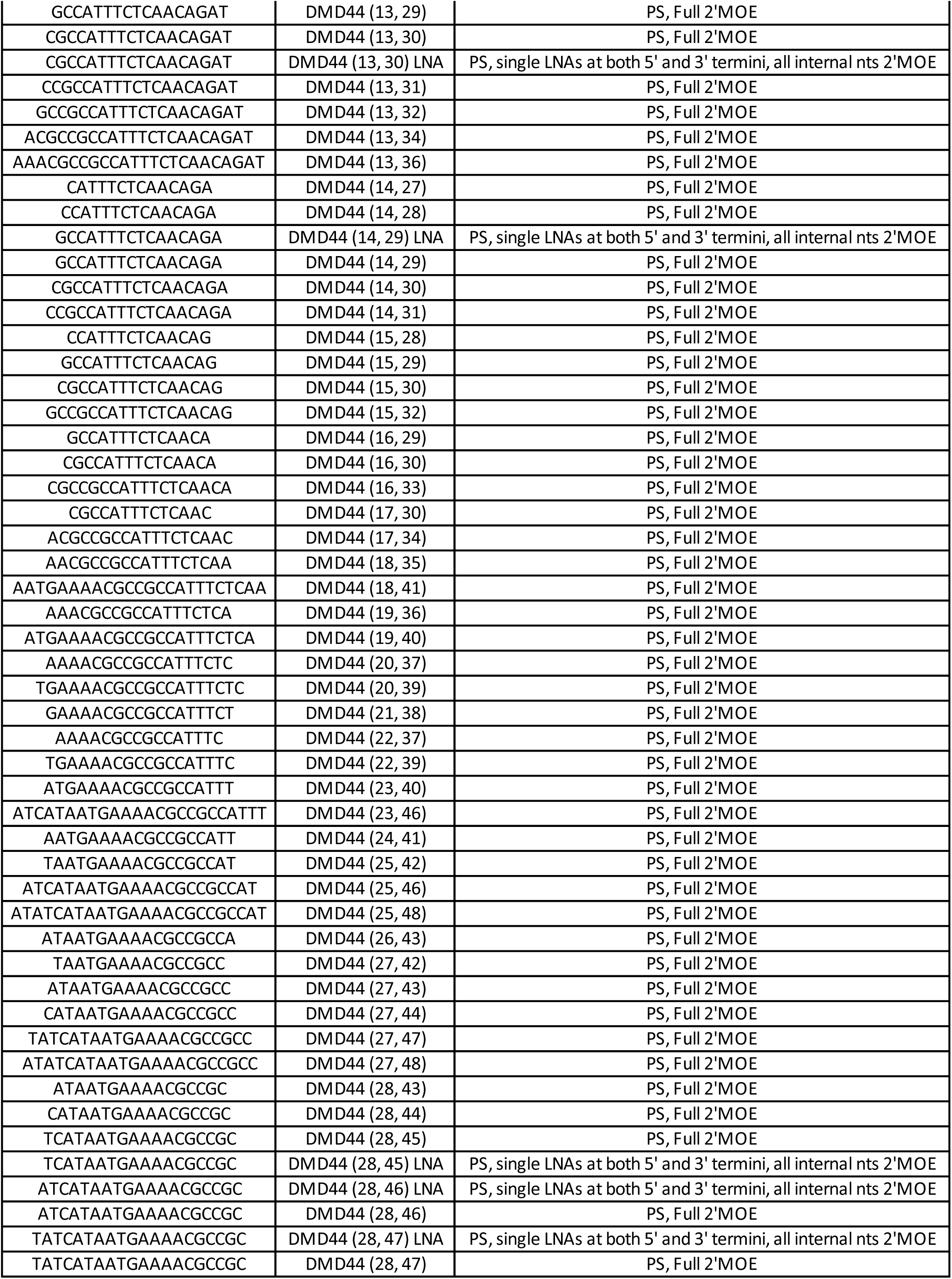
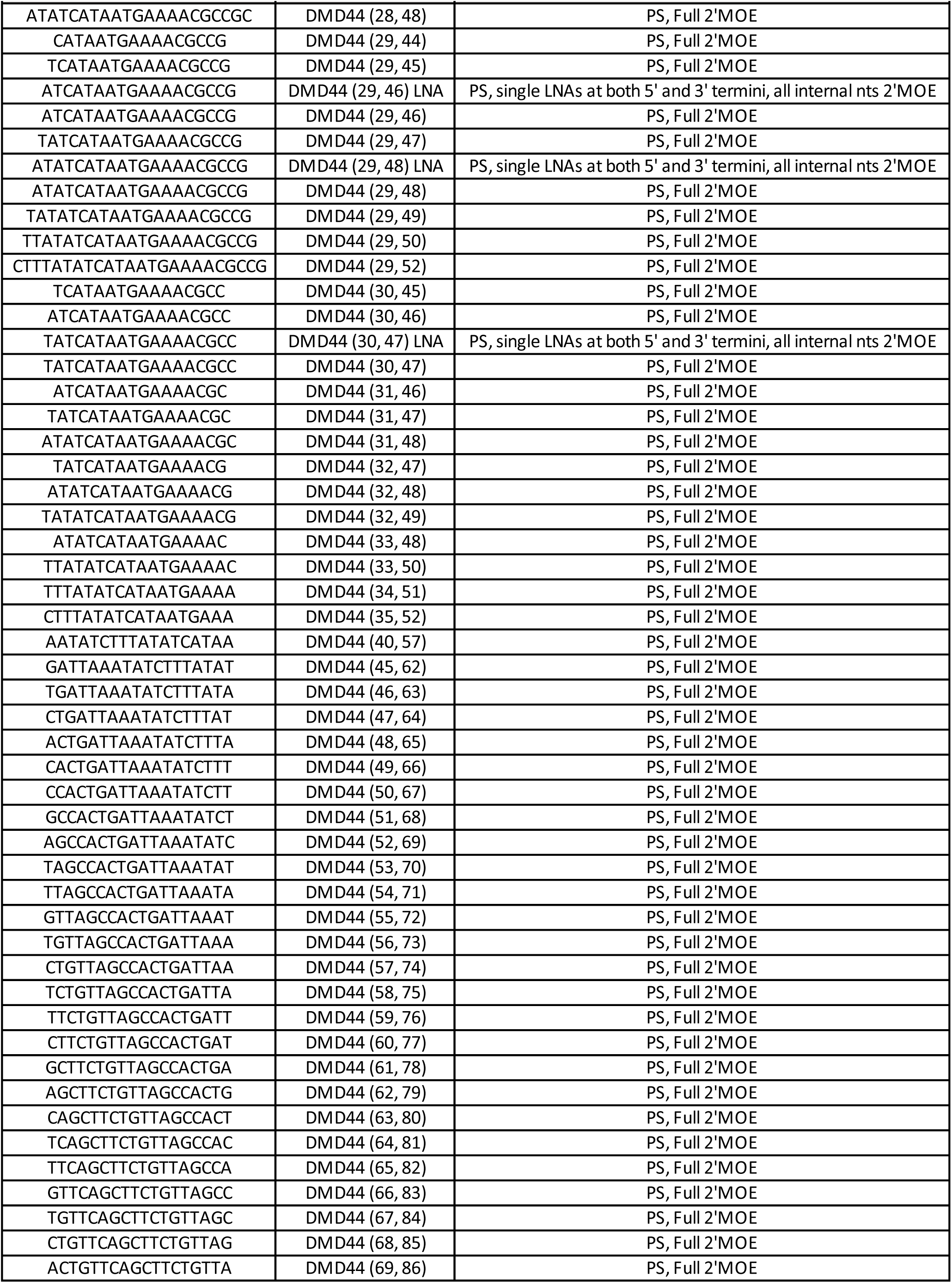
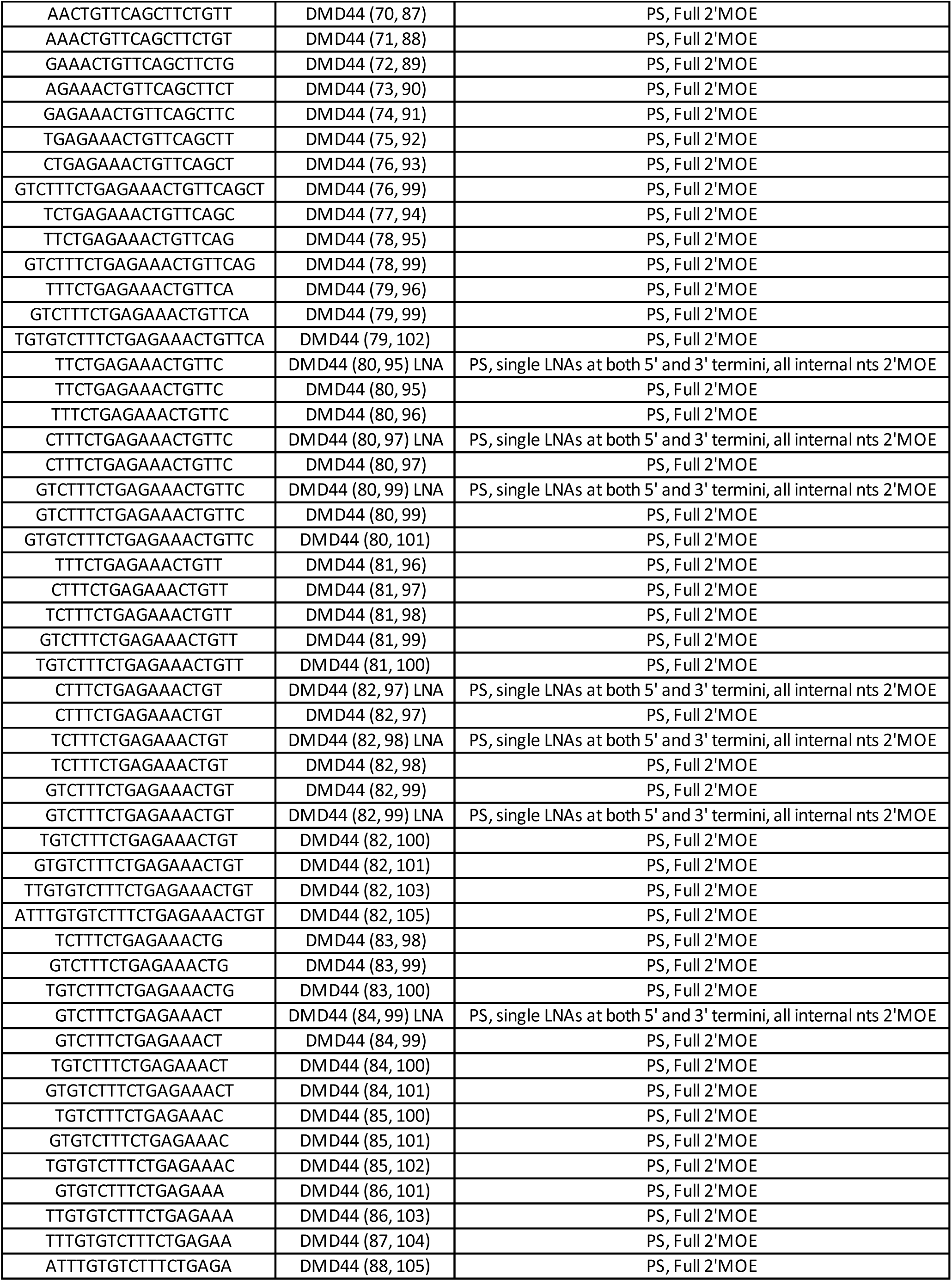
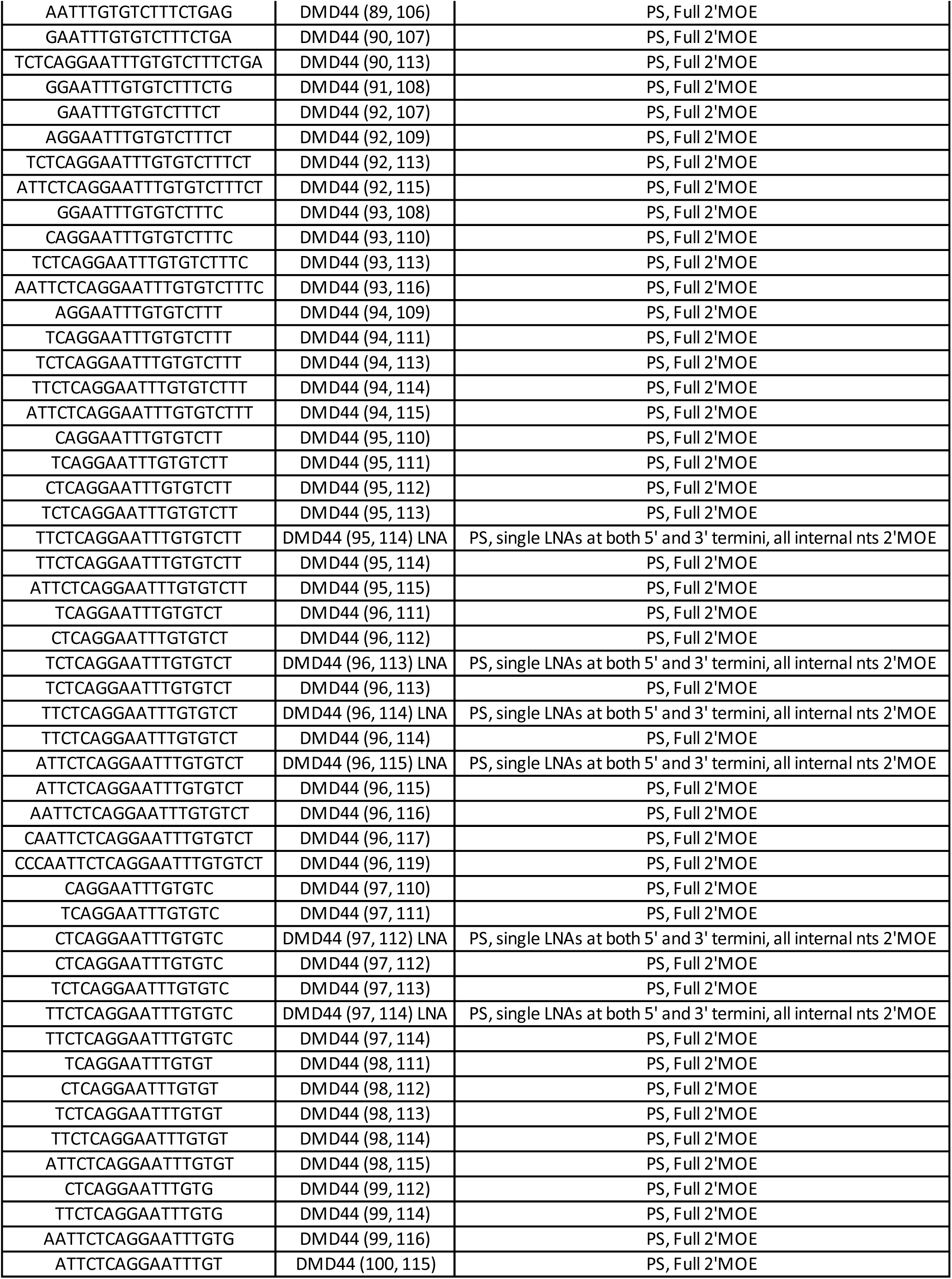
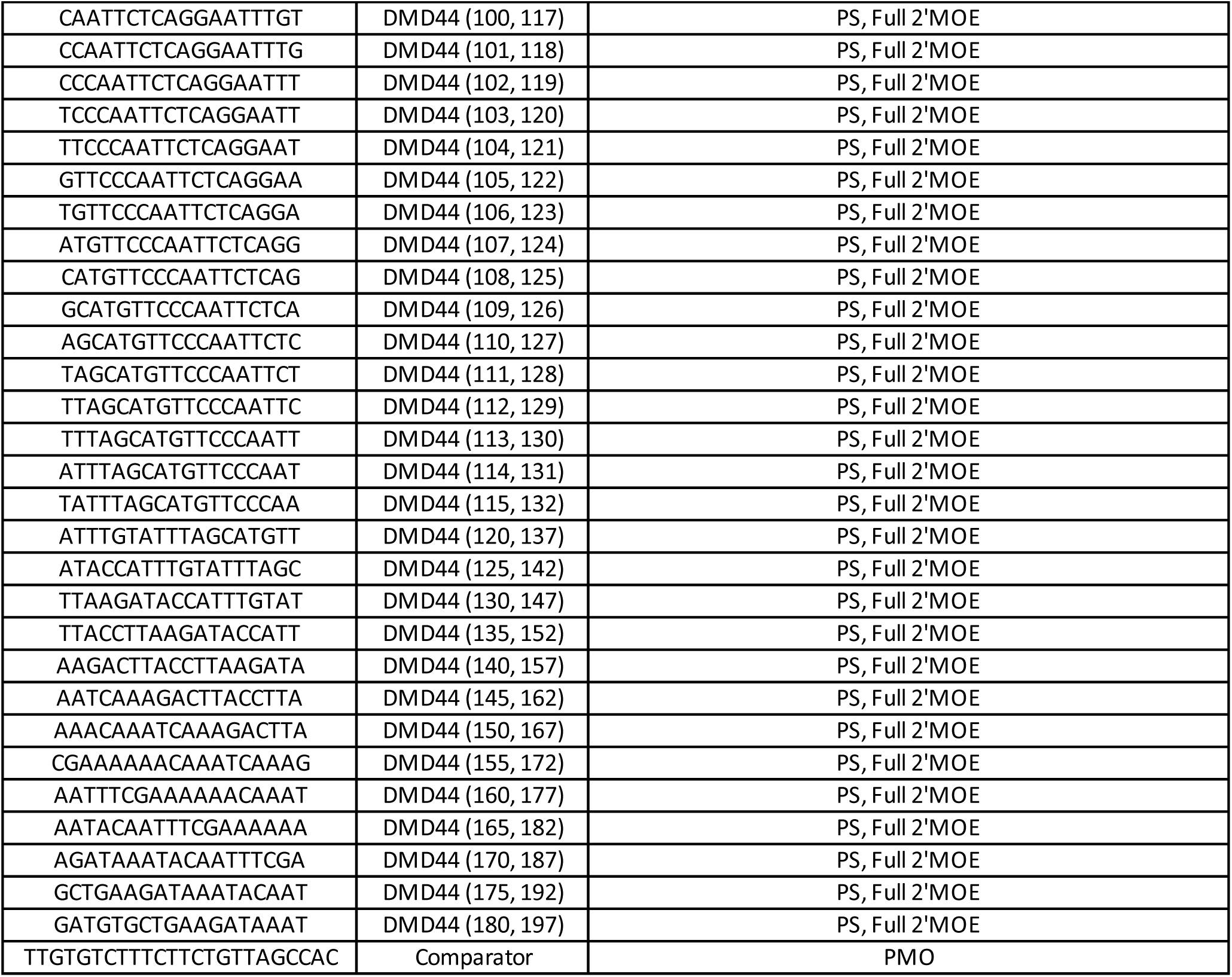

**Figure S1.**
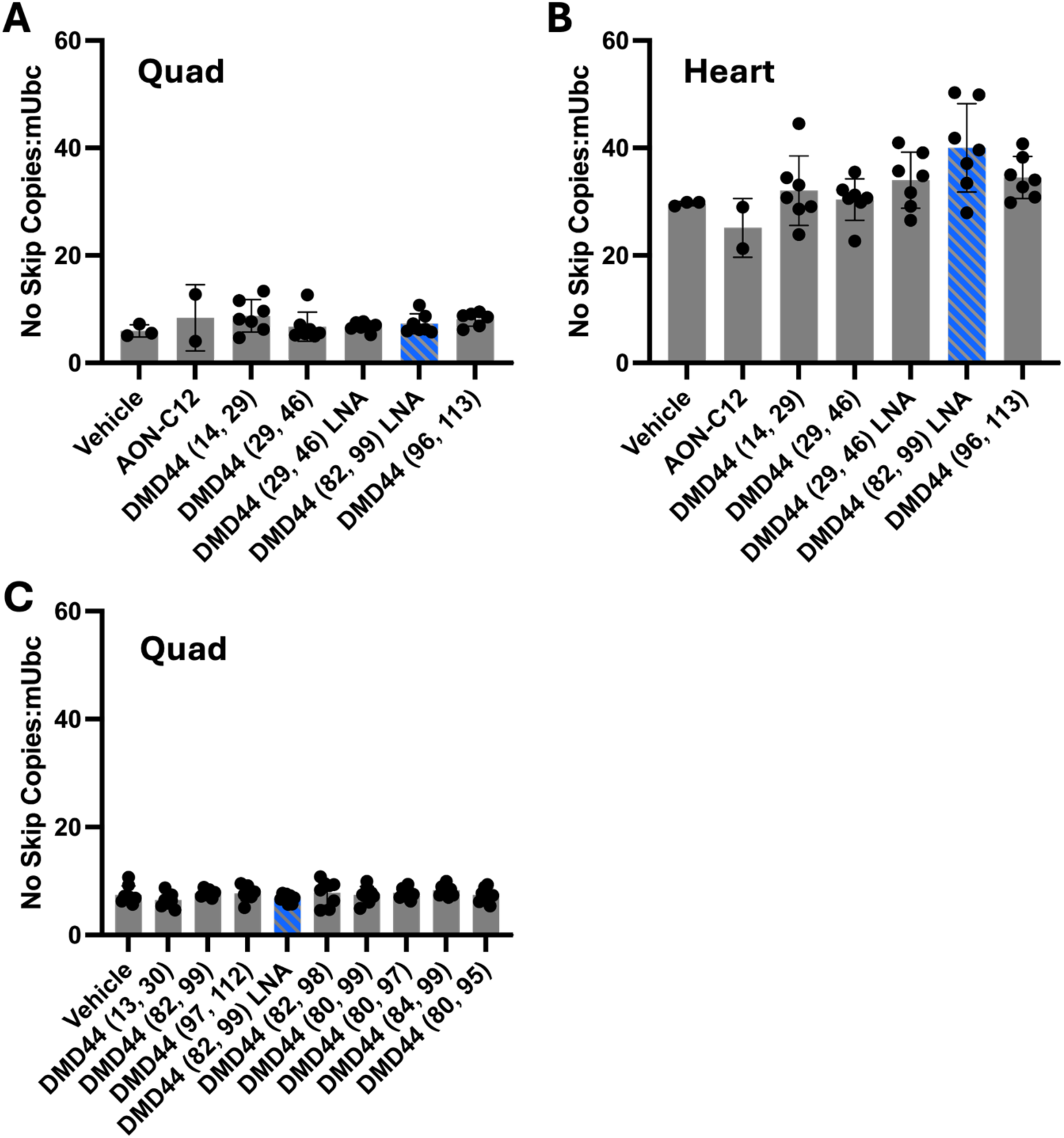
None of the Exon 44 skipping ASOs impacted levels of non-skipped dystrophin mRNA in hDMD mice. In “study 1”, where a set of ASOs were tested in the hDMD mice at 1.71 µmol/kg, none impacted the levels of the non-skipped dystrophin mRNA relative to vehicle treatment in Quad (A) and Heart (B). Similarly, no ASOs at 3.42 µmol/kg impacted non-skipped dystrophin mRNA levels relative to vehicle treatment in Quad.

**Figure S2.**
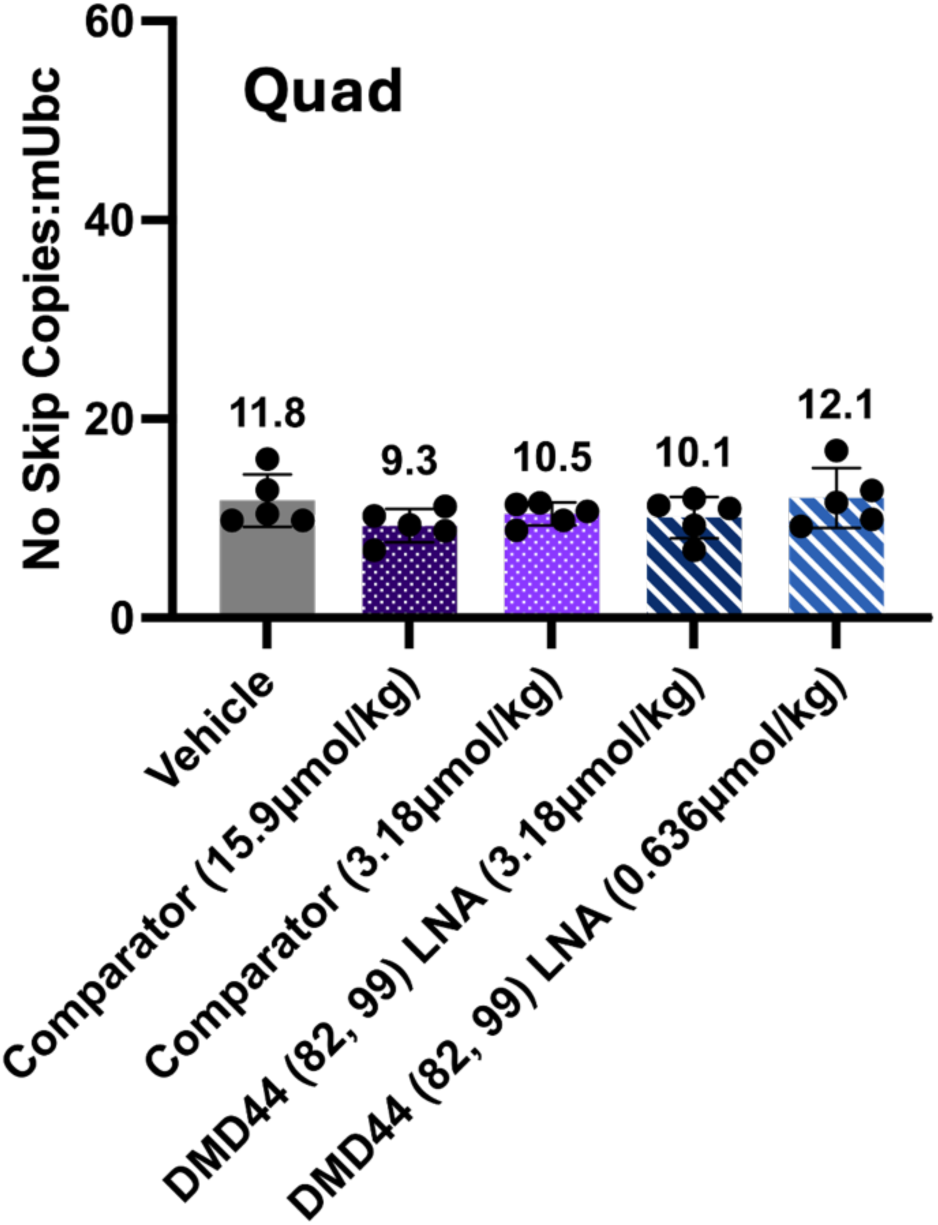
None of the Exon 44 skipping ASOs impacted levels of non-skipped dystrophin mRNA in hDMDdel45/mdx mice. In study 2, where a set of ASOs were tested in the hDMDdel45/mdx mice, none impacted the levels of the non-skipped dystrophin mRNA relative to vehicle treatment in Quad.

